# Dynamics of RNA polymerase II and elongation factor Spt4/5 recruitment during activator-dependent transcription

**DOI:** 10.1101/2020.06.01.127969

**Authors:** Grace A. Rosen, Inwha Baek, Larry J. Friedman, Yoo Jin Joo, Stephen Buratowski, Jeff Gelles

## Abstract

In eukaryotes, RNA polymerase II (RNApII) transcribes messenger RNA from template DNA. Decades of experiments have identified the proteins needed for transcription activation, initiation complex assembly, and productive elongation. However, the dynamics of recruitment of these proteins to transcription complexes, and of the transitions between these steps, are poorly understood. We used multi-wavelength single-molecule fluorescence microscopy to directly image and quantitate these dynamics in a budding yeast nuclear extract that reconstitutes activator-dependent transcription in vitro. A strong activator (Gal4-VP16) greatly stimulated reversible binding of individual RNApII molecules to template DNA, with no detectable involvement of RNApII-containing condensates. Binding of labeled elongation factor Spt4/5 to DNA typically followed RNApII binding, was NTP-dependent, and was correlated with association of mRNA-binding protein Hek2, demonstrating specificity of Spt4/5 binding to elongation complexes. Quantitative kinetic modeling shows that only a fraction of RNApII binding events are productive and implies a rate-limiting step, probably associated with recruitment of general transcription factors, needed to assemble a transcription-competent pre-initiation complex at the promoter. Spt4/5 association with transcription complexes was slowly reversible, with DNA-bound RNApII molecules sometimes binding and releasing Spt4/5 multiple times. The average Spt4/5 residence time was of similar magnitude to the time required to transcribe an average length yeast gene. These dynamics suggest that a single Spt4/5 molecule remains associated during a typical transcription event, yet can dissociate from RNApII to allow disassembly of abnormally long-lived (i.e., stalled) elongation complexes.

**Significance Statement:** The synthesis of a eukaryotic messenger RNA molecule involves the association of RNA polymerase and dozens of accessory proteins on DNA. We used differently colored fluorescent dyes to tag DNA, RNA polymerase II, and the elongation factor Spt4/5 in yeast nuclear extract, and then observed the assembly and dynamics of individual molecules of the proteins with single DNA molecules by microscopy. The observations quantitatively define an overall pathway by which transcription complexes form and evolve in response to an activator protein. They suggest how molecular complex dynamics may be tuned to optimize efficient RNA production.

## Introduction

As the initiating event in eukaryotic gene expression, mRNA transcription by RNA polymerase II (RNApII) is regulated at several steps (1–3). First, gene-specific transcription activators recognize specific DNA binding sites upstream of the transcription start site. Once bound, these activators recruit histone acetyltransferases and chromatin remodelers to move promoter-proximal nucleosomes, making the DNA accessible for RNApII pre-initiation complex (PIC) formation. Activators further accelerate pre-initiation complex (PIC) formation by recruiting the basal initiation factors via contacts with the Mediator/RNApII complex and TFIID. Once assembled, the PIC undergoes several ATP-dependent rearrangements, including promoter melting and phosphorylation of the RNApII largest subunit Rpb1. Phosphorylation of the Rpb1 C-terminal domain (CTD) releases Mediator, facilitating conversion to a transcription elongation complex (EC). During elongation, a series of CTD kinases and phosphatases creates dynamic phosphorylation patterns that differentially recruit mRNA processing and chromatin modifying enzymes to early and late stages of transcription. Transcription can be regulated at the level of PIC assembly, by controlling the transition to elongation, or by early termination events that attenuate gene expression.

The PIC to EC transition requires the exchange of initiation and elongation factors that contact various surfaces of RNApII. For example, structural studies show that the elongation factor Spt4/5 (also known as DRB-sensitivity inducing sensitivity factor, or DSIF, in metazoans) and the initiation factor TFIIE occupy overlapping space on RNApII. The archaeal Spt5 and TFE (TFIIE) homologs compete for binding to the RNAP clamp, a mobile domain that regulates template DNA movement into or out of the polymerase active site cleft (4). Similarly, the TFIIF initiation factor and Paf1 complex elongation factor may also bind RNApII mutually exclusively (5).

Eukaryotic Spt5 consists of a NusG N-terminal (NGN) domain, three to five C-terminal Kyprides-Ouzounis-Woese (KOW) domains, and a C-terminal region (CTR) that is phosphorylated during transcription elongation (6). In a manner analogous to its bacterial homolog NusG, the Spt5 NGN and KOW1 domains bridge the gap across the RNApII cleft in the EC (5). Spt4, although non-essential for viability in yeast and not conserved in bacteria, dimerizes with the NGN domain to stabilize this interaction. A large body of data shows that Spt4/5 acts to positively regulate elongation (6). Both bacterial and yeast Spt5 homologs increase RNApII elongation rate by reducing transcriptional pausing (6). Human Spt4/5 overcomes promoter-proximal pausing induced by the Negative Elongation Factor, NELF (7). Chromatin immunoprecipitation experiments in *Saccharomyces cerevisiae* show decreased RNApII density at the 3′ end of genes in strains mutated in Spt4/5 (8–11). Stimulation of elongation by Spt4/5 may involve multiple mechanisms, including stabilization of a more processive RNApII conformation, steric blockage of template release, interaction between KOW domains and the RNA transcript (12, 13), and recruitment of additional elongation factors to the KOW and CTR domains (6, 14–16).

To study the factors that drive transcription activation, PIC formation, and transition to elongation, we have used quantitative mass spectrometry to analyze transcription complexes in vitro. Complexes were assembled on an immobilized DNA transcription template in yeast nuclear extract (17). This system uses a model promoter consisting of the *CYC1* core promoter driven by five upstream Gal4 binding sites. We used the synthetic protein Gal4-VP16, which fuses the Gal4 DNA-binding domain to the herpes simplex virus VP16 activation domain, resulting in a powerful transcription activator in both mammalian (18) and yeast systems (19). We previously showed that the Gal4 DNA-binding domain, when fused to either VP16 or the native yeast Gcn4 activation domain, strongly recruits the NuA4 and SAGA histone acetyltransferases, the Swi/Snf remodeling complex, and Mediator and other PIC components, to both naked DNA and chromatinized templates (17). Furthermore, this system faithfully recapitulates the transition to elongation and the exchange of factors mediated by the Rpb1 CTD phosphorylation cycle (20). While this system has proven useful for studying dynamics, the time resolution of the mass spectrometry experiments was on the order of minutes rather than seconds. Furthermore, since the transcription complexes were analyzed in bulk, results represent the averaged behaviors of the molecular ensemble, yielding only incomplete information about the pathways of complex assembly.

To provide increased time resolution and the ability to monitor reaction pathways of activated transcription events on individual DNA molecules, here we adapt the immobilized template assay for single-molecule light microscopy. Colocalization Single Molecule Spectroscopy (CoSMoS) is a multi-wavelength single-molecule fluorescence method well suited for characterizing molecular mechanisms (21, 22). We have previously used this approach to analyze the pathways of molecular complex formation and function in bacterial transcription and pre-mRNA splicing (22–29). By fluorescently labeling the endogenous RNApII and Spt4/5 in nuclear extract, as well as the DNA template, with three differently colored dyes, we can monitor the arrival and departure of the individual protein molecules from the template DNA. Because these experiments are done in nuclear extracts, they complement other single molecule studies of RNApII transcription that used a limited set of purified factors (30). As the full complement of nuclear proteins is present, the experimental system can recapitulate a more complete set of transcription processes and thus provide a better approximation of events in vivo. Conversely, the approach overcomes some of the technical challenges of single-molecule studies of transcription in live cells (31), allowing more complete kinetic analysis of multiple components associating simultaneously with an isolated DNA molecule leading up to and during the synthesis of a single RNA molecule.

Our experiments, and the numerical modeling of the dynamics they reveal, quantitatively define a minimal kinetic mechanism for activator-dependent transcription in this system. The analysis distinguishes productive and non-productive RNApII binding and defines key rate-limiting steps in initiation. Furthermore, the experiments reveal reversible association of Spt4/5 with transcription elongation complexes and suggest that these dynamics are tuned to the timescale of yeast gene transcription.

## Results

### A system for imaging RNApII and Spt4/5 binding to promoter DNA

To detect transcription complex formation by single-molecule fluorescence, we constructed a *S. cerevisiae* strain in which the Rpb1 subunit of RNApII is fused to a C-terminal SNAP tag (32) and the Spt5 subunit of Spt4/5 is fused to a C-terminal dihydrofolate reductase (DHFR) tag (33). The tagged proteins are expressed at levels similar to the corresponding untagged proteins in the parental strain, and the tags did not perturb yeast cell growth rate or transcription activity in nuclear extracts when compared to the parental strain (**Fig. S1**). Rpb1-SNAP in the extract was labeled with SNAP-Surface 549, a SNAP substrate conjugated to the green-excited dye DY549. Imaged transcription reactions also included Cy5-trimethoprim (Cy5-TMP) to non-covalently label the DHFR-tagged Spt5 subunit of Spt4/5 with the red-excited dye Cy5. We transcribed a DNA template containing five Gal4 binding sites upstream of the *CYC1* core promoter, which drives a downstream 300 bp cassette encoding a G-less RNA (**Fig. 1A**). On supercoiled (**Fig. S1C, D**) or linear templates (20), robust transcription was seen upon addition of the transcription activator Gal4-VP16 (18) and nucleotides (NTPs) ATP, UTP, CTP, and 3′ O-methyl-GTP. The omission of GTP suppresses production of non-specific transcripts initiating from cryptic promoters and DNA ends.

**Figure 1.**
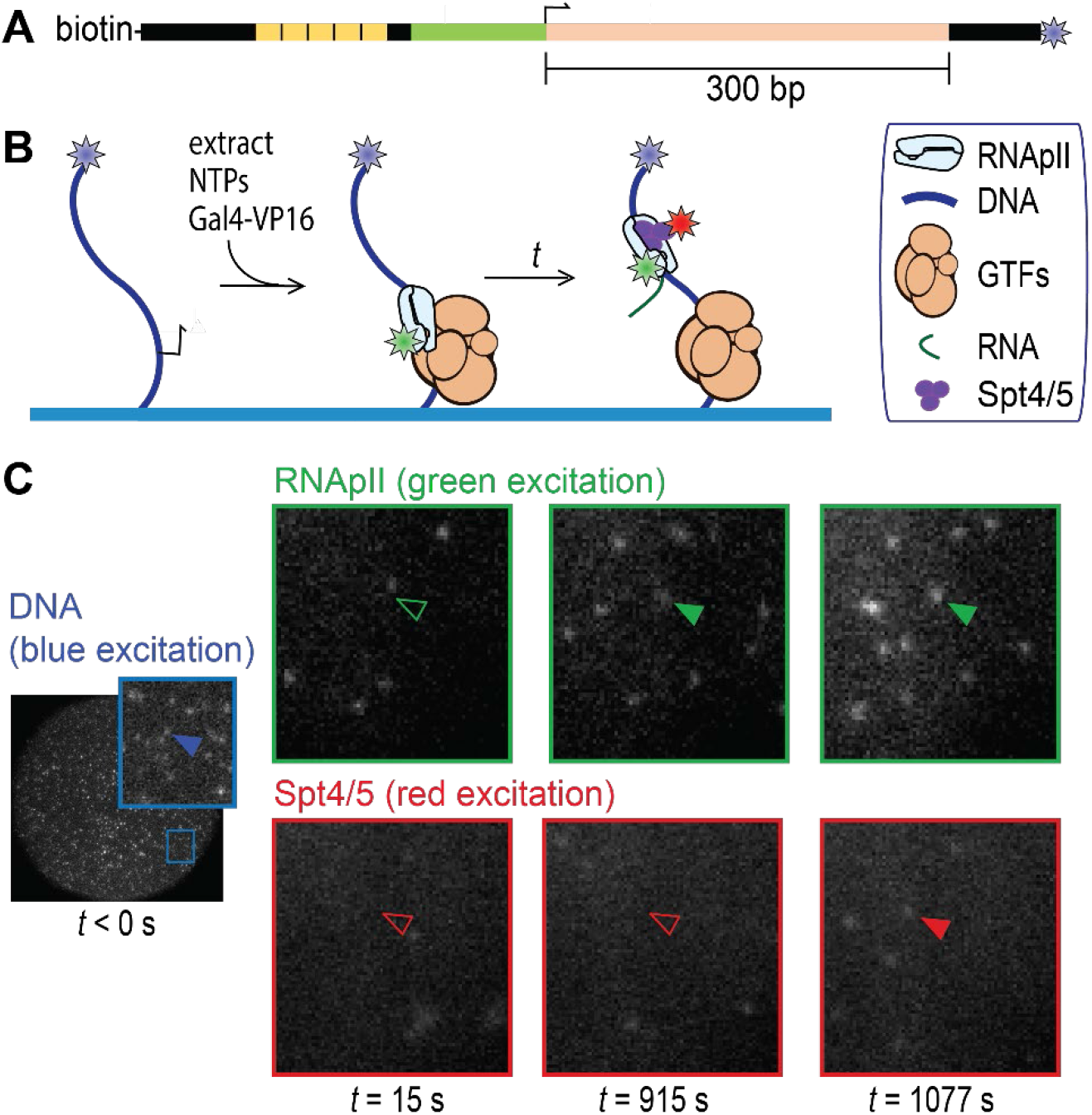
Detection of RNApII and Spt4/5 binding to individual surface-tethered DNA^488^ molecules in Rpb1^SNAP549^/Spt5^DHFR-Cy5^ *S. cerevisiae* nuclear extract. **(A)** Schematic of DNA^488^ transcription template. This DNA contains five upstream Gal4 binding sites (yellow) and the *CYC1* core promoter (green) with its transcription start site (bent arrow), followed by a 300 bp cassette (pink) encoding a G-less RNA. The template has attached biotin and AF488 dye (blue star) moieties. **(B)** Experimental scheme. DNA^488^ molecules immobilized on the surface of a flow chamber (blue) were at time *t* = 0 incubated with yeast nuclear extract containing dye (stars) -labeled proteins Rpb1^SNAP549^ and Spt5^DHFR-Cy5^ along with unlabeled general transcription factors (GTFs) and other nuclear proteins, and which was supplemented with recombinant Gal4-VP16 activator. RNApII and Spt4/5 binding to DNA were detected as colocalization of spots of green- and red-excited fluorescence at locations of blue-excited DNA spots. **(C)** Images of the same microscope field of view (65 × 65 μm) in the red-, green-, and blue-excited fluorescence channels taken at various times before (blue) and after (red and green) extract addition at time *t* = 0. Insets show magnified views of the marked region. Absence or presence of a spot of fluorescence colocalized with a particular DNA molecule are shown by open and filled arrowheads, respectively.

To visualize RNApII and Spt4/5 association with individual DNA molecules, transcription templates were PCR amplified using an upstream oligonucleotide labeled with biotin and downstream oligo carrying a blue-excited dye (**Fig. 1A**). The templates were tethered to the surface of a glass flow chamber and then observed by micromirror multi-wavelength TIRF microscopy (21). DNA locations were mapped in the blue channel (DNA^488^), as were a randomly selected control set of “no-DNA” spots used to monitor background binding. Labeled extract, Gal4-VP16, and NTPs (ATP, UTP, CTP, and 3’-O-MeGTP) were then added to begin the reaction (**Fig. 1B**). Candidate binding events for RNApII and/or Spt4/5 were detected in the green (Rpb1^SNAP549^) and red (Spt5^DHFR-Cy5^) fluorescence channels, respectively (**Fig. 1C**).

### Activator dependence of RNApII recruitment

Time records of Rpb1^SNAP549^ fluorescence at individual DNA locations showed association of single or sometimes two or more RNApII molecules with the template (**Fig. 2A**). Of *N* = 331 DNA^488^ molecules observed in a representative experiment, 294 (89%) exhibited binding of at least one Rpb1^SNAP549^ molecule over the course of the 40 min recording. Control no-DNA locations showed far fewer Rpb1^SNAP549^ binding events (**Fig. 2A, B**), confirming that most binding is DNA-specific.

**Figure 2:**
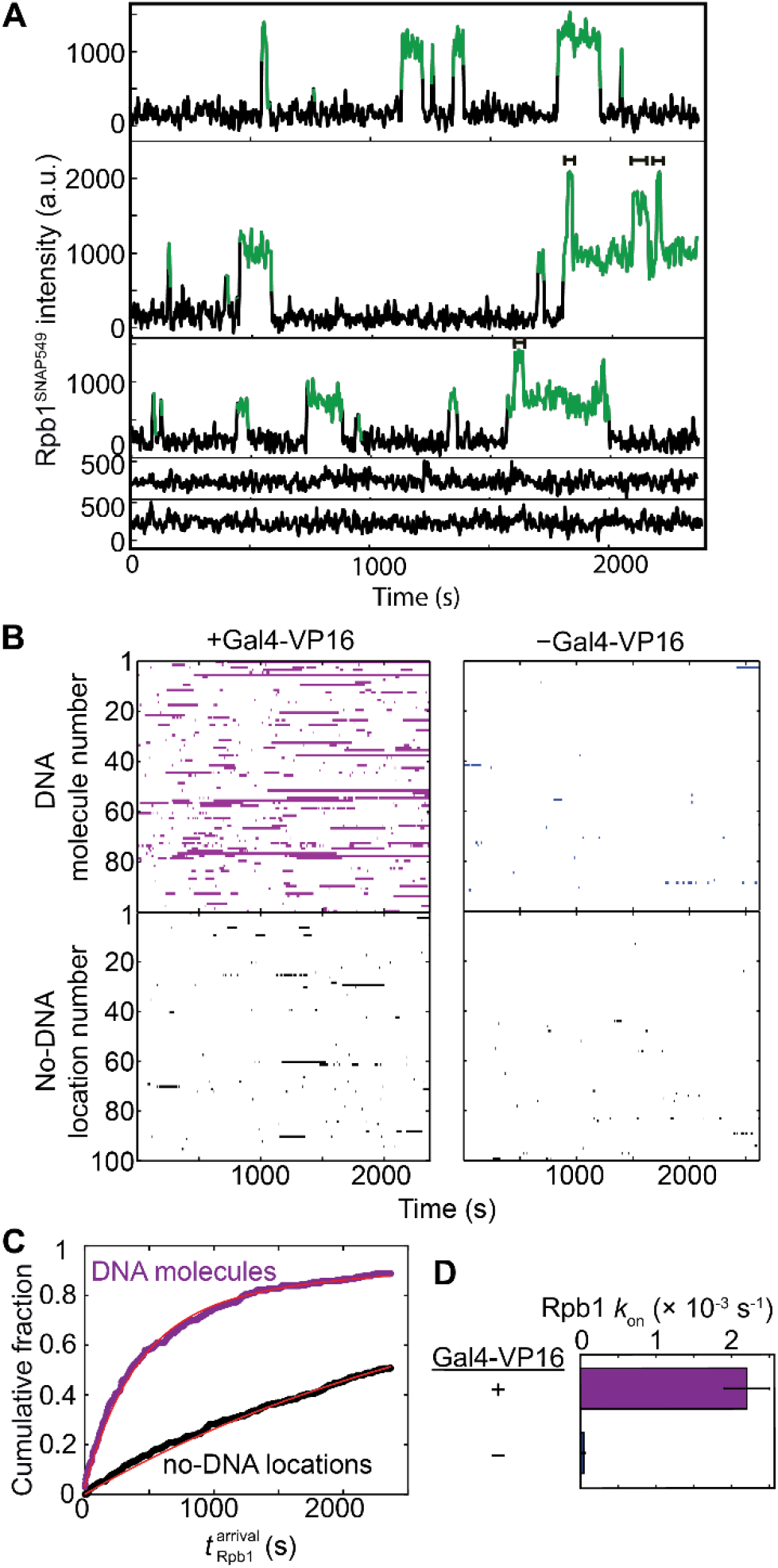
Kinetics of RNApII association with promoter DNA in the experiment shown in **Fig. 1**. **(A)** Example time records showing Rpb1^SNAP549^ fluorescence colocalized at the positions of three different single DNA^488^ molecules (top three records) and two control locations without visible DNA^488^ molecules (bottom two). Green points designate times at which there was a colocalized spot of Rpb1^SNAP549^ fluorescence as detected by an objective spot-recognition algorithm (59). Brackets mark time intervals during which more than one Rpb1^SNAP549^ molecule was present. **(B)** Rastergrams of Rpb1^SNAP549^ colocalization at the locations of 100 randomly selected DNA^488^ molecules (top panels) and 100 randomly selected control no-DNA locations (bottom panels) from the same recording. Each horizontal line of the plot is data from a single location showing times with (colored bars) or without (white space) a colocalized Rpb1^SNAP549^ fluorescent spot. Data are from the experiment of **Fig. 1** (purple) plus a negative control experiment in which no Gal4-VP16 activator was added (blue). **(C)** Cumulative distribution of the time intervals prior to the first Rpb1^SNAP549^ association seen on each DNA^488^ molecule (purple) and at no-DNA^488^ control locations (black) in the presence of Gal4-VP16, along with fits (red) to an exponential binding model (a single rate-limiting step). Parameters of this and analogous fits to the negative control and the numbers of observations for all fits are given in **Table S1**. **(D)** Apparent first-order rate constants (± S.E.) for Rpb1 association with template DNA derived from the model fits.

Measurement of the delay time from the addition of extract to first observed binding Rpb1^SNAP549^ on each DNA^488^ molecule revealed an exponential distribution with an apparent first-order association rate constant of (2.2 ± 0.3) × 10^−3^ s^−1^ (**Fig. 2C, D**). This number usually was within a factor of ~2.5 in experiments using different extracts, including those where Rpb1 was tagged with DHFR instead of SNAP, suggesting that extract batch and tag identity did not greatly alter binding kinetics (**Table S1**). These results are consistent with initial RNApII association being dominated by a single rate-limiting step under these experimental conditions. The binding rate constant was reduced by more than an order of magnitude in the absence of Gal4-VP16 (**Fig. 2B, D**, blue), demonstrating that the vast majority of RNApII DNA binding detected in this system is activator-dependent.

The mean lifetime for RNApII occupancy of DNA in the presence of NTPs and activator was 79 ± 2 s (S.E.; *N* = 1,259), although some individual Rpb1 molecules remained associated with DNA for hundreds of seconds (**Fig. 2A, 2B**). However, multiple molecules of Rpb1^SNAP549^ were sometimes observed simultaneously on an individual DNA (**Fig. 2A**), so the average lifetime of individual molecules may be shorter. Multiple RNApII molecules may reflect simultaneous presence of initiation and elongation complexes. When NTPs were absent, we still observed significant RNApII association with DNA; a similar result was obtained in the presence of NTPs when the inhibitor α-amanitin was added (**Fig. S2; Table S1**). These results are consistent with the ability of RNApII to participate in pre-initiation complexes under these conditions, even though NTP depletion or α-amanitin both block productive transcription.

### Spt4/5 binding is specific to elongation complexes

In the same experiment monitoring Rpb1^SNAP549^, the arrival of Spt4/5 was visualized by colocalization of Spt5^DHFR-Cy5^ with the DNA^488^ template molecules (**Fig. 1, Fig. 3A, Fig. S3**). Neither RNApII nor Spt4/5 kinetics were affected by tagging of the other protein (**Table S1; Table S2**), suggesting that the tagged proteins behave similarly to wild-type.

**Figure 3.**
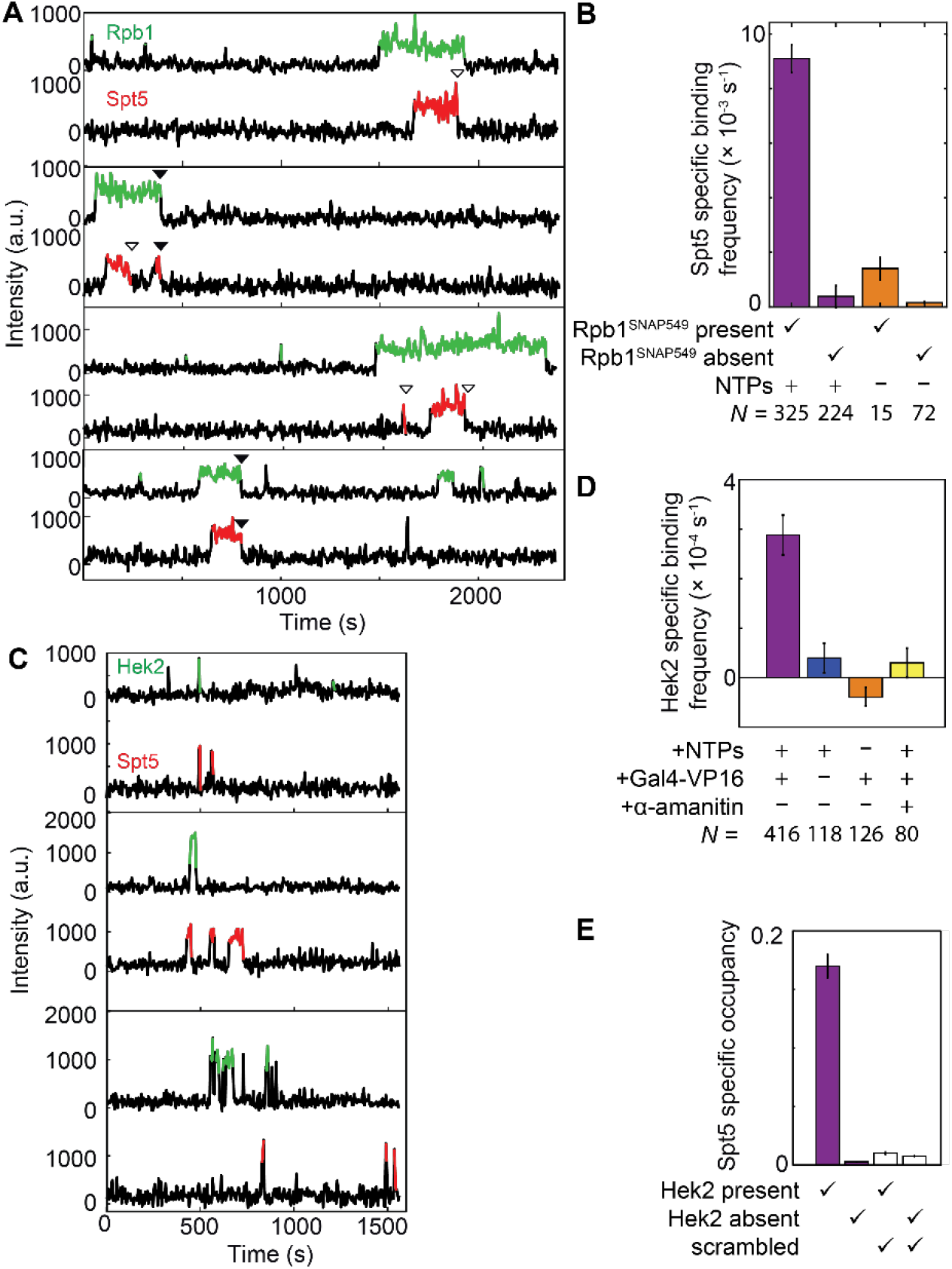
Correlation of Spt4/5 binding at individual DNA molecules with binding of RNApII **(A, B)** or Hek2 **(C – E)**. **(A)** Example time records of Rpb1^SNAP549^ and Spt5^DHFR-Cy5^ fluorescence at four individual DNA^488^ locations, taken from the Rpb1^SNAP549^/Spt5^DHFR-Cy5^ experiment shown in **Fig. 1**. Colored intervals indicate times at which a fluorescent Rpb1^SNAP549^ (green) and/or Spt5^DHFR-Cy5^ (red) spot colocalized to the DNA^488^ molecule being monitored. Each Spt5^DHFR-Cy5^ departure is marked according to whether it occurred before (open triangles) or simultaneously with (closed triangles) Rpb1^SNAP549^ departure. Additional example records are shown in **Fig. S3**. **(B)** DNA-specific binding frequencies (± S.E.) of Spt5^DHFR-Cy5^, calculated separately for time intervals when Rpb1^SNAP549^ was present or absent at the DNA. DNA-specific binding frequency is calculated as the frequency seen at DNA locations that is in excess of the background non-specific binding seen at no-DNA locations. **(C)** Examples of time records of Hek2^SNAP549^ and Spt5^DHFR-Cy5^ fluorescence over time at three individual DNA^488^ locations taken from an experiment using a dual-tagged Hek2^SNAP549^/Spt5^DHFR-Cy5^ yeast nuclear extract in the presence of NTPs and Gal4-VP16. **(D)** Frequencies (± S.E.) of DNA-specific Hek2^SNAP549^ binding under transcription conditions (purple) and in negative controls (blue, orange, and yellow); see **Table S3**. **(E)** Mean fraction (± S.E.M.) of time that a DNA^488^ molecule had a colocalized Spt5^DHFR-Cy5^, after correction for non-specific surface binding. Separate analyses were conducted for time points at which Hek2^SNAP549^ was present or was absent (**Table S4)**. Data may underestimate the true occupancy of DNA locations by Spt4/5 due to possible incomplete labeling of Spt5^DHFR-Cy5^, but this factor is constant across all four bars. Open bars show results of a scrambled negative control analysis of the same data (see Methods) showing that the correlation of Spt5 binding with Hek2 binding is not a statistical artifact. Data in (D, purple bar) and (E) are aggregated from the experiment in (C) and one additional replicate.

Spt4/5 binding at the location of a particular DNA molecule almost always occurred while RNApII was detected on that DNA (**Fig. 3A, Fig. S3**). In particular, Spt5^DHFR-Cy5^ DNA-specific binding (i.e., binding in excess of the background seen at no-DNA locations) occurred >10-fold more frequently when an Rpb1^SNAP549^ spot was present than when absent (**Fig. 3B**, purple bars). The rare occurrences of Spt5^DHFR-Cy5^ binding without Rpb1^SNAP549^ on the DNA (33 of 549 total Spt5^DHFR-Cy5^ binding events) may reflect non-specific binding to the slide surface or specific binding to unlabeled or photobleached Rpb1^SNAP549^. Based on these observations, at least 94 ± 1 % of the RNApII molecules in the extract were fluorescent, consistent with observations that dye labeling of Rpb1^SNAP^ was saturated under the labeling conditions used (**Fig. S4**). Even though Spt4/5 contacts both RNApII protein and template DNA in ECs (13, 34), the colocalization data suggest that Spt4/5 recruitment to DNA requires the presence of RNApII. In agreement with this idea, we almost invariably (288 of 312 instances) saw that RNApII arrived on DNA before Spt4/5 when complexes that contained both proteins were formed.

Consistent with a direct interaction between Spt4/5 and RNApII, 42 out of 312 co-localized Rpb1^SNAP549^/Spt5^DHFR-Cy5^ spots departed from the DNA simultaneously, within experimental time resolution (i.e., within ±1 video frame; **Fig. 3A**, filled arrows), as expected for an RNApII•Spt4/5 complex dissociating from the DNA as a unit. In most other events (241 of the 270 co-localized Rpb1^SNAP549^ and Spt5^DHFR-Cy5^ spots without simultaneous departure), Spt4/5 departed before RNApII dissociation. Based on all these results, we conclude that most (and possibly all, if the exceptions reflect Rpb1^SNAP549^ photobleaching) Spt4/5 binding is dependent on the presence of DNA-bound RNApII.

Some key Spt4/5 contacts with RNApII seen in EC structural models appear prohibited in the PIC due to obstruction by TFIIE or other PIC components (13, 35, 36), and multiple observations suggest that Spt4/5 associates stably with ECs but not PICs (20, 37). Consistent with this suggestion, we saw that the frequency with which Spt5^DHFR-Cy5^ associated with a DNA containing Rpb1^SNAP549^ was >6-fold higher in the presence of added NTPs than in NTP depletion conditions in which ECs cannot form [(9.4 ± 0.5) ×10^−3^ s^−1^ vs. (1.4 ± 0.1) ×10^−3^ s^−1^, respectively; **Fig. 3B**]. Simultaneous occupancy of a DNA by multiple Spt5^DHFR-Cy5^ molecules was almost never seen (2 out of 331 DNA molecules examined). Taken together, these data suggest that Spt4/5 is interacting with ECs and that at most one EC is typically present on each DNA in these experiments.

As an independent test for Spt5 binding to ECs, we used an Spt5^DHFR-Cy5^/Hek2^SNAP549^ nuclear extract to monitor colocalization of Spt5 with Hek2, an RNA transcript-binding protein known to associate with elongation complexes generated in the nuclear extract system used here (20, 38). In contrast to Spt5^DHFR-Cy5^, Hek2^SNAP549^ association with surface-tethered DNA locations was much shorter-lived (**Fig. 3C**), possibly due to low affinity or competition by other RNA-binding proteins for nascent transcript binding. Nevertheless, Hek2^SNAP549^ colocalization with DNA was transcription-dependent, as no significant binding over background was detected in controls lacking activator or NTPs or containing α-amanitin (**Fig. 3D**). Thus, we propose that DNA-specific presence of Hek2^SNAP549^ can serve as a proxy for detection of an EC-associated transcript. Consistent with Spt4/5 binding being restricted to ECs, the probability of Spt5^DHFR-Cy5^ being present simultaneously with Hek2^SNAP549^ was more than 10-fold higher than that for Spt5^DHFR-Cy5^ presence during intervals when Hek2^SNAP549^ was absent (**Fig. 3E**). To exclude the possibility that the apparent association between Spt4/5 and Hek2 presence resulted from coincidental independent binding events, the Spt5^DHFR-Cy5^ and Hek2^SNAP549^ occupancy records from different individual DNAs molecules were randomly paired and the analysis repeated (“scrambled” in **Fig. 3E** and **Table S4**). The correlation was completely lost, confirming that the observed Hek2 and Spt4/5 colocalization represents binding to the same molecular complex. Thus, the Hek2 data provides additional support that the observed Spt4/5 binding is specific for ECs.

### Dynamics of initiation and elongation complex formation

Despite most DNAs having one or more Rpb1^SNAP549^ binding events (**Fig. 2C**; **Fig. 4A**, purple), only a subpopulation of these events showed colocalization with Spt5^DHFR-Cy5^ (examples are marked ‘p’ in **Fig. S3**). As discussed above, the exponential distribution of initial Rpb1^SNAP549^ binding times (**Fig. 2C**) indicates a single rate-limiting step for association. Strikingly, this was not the case for times to the first *productive* RNApII associations (defined as the first Rpb1^SNAP549^ binding event during which Spt5^DHFR-Cy5^ was also colocalized; **Fig. 4A**, turquoise). These productive events exhibited a non-exponential distribution that increases steeply only after a distinct lag time of ~100 s. Such a distribution is inconsistent with a simple model in which all RNApII associations have the same chance of forming an Spt4/5-containing EC. Instead, they suggest an additional slow event must occur during incubation of DNA template with extract to allow elongation complex production. In such a situation, later-binding RNApII molecules are more likely to produce elongation complexes, resulting in the observed lag kinetics.

**Figure 4.**
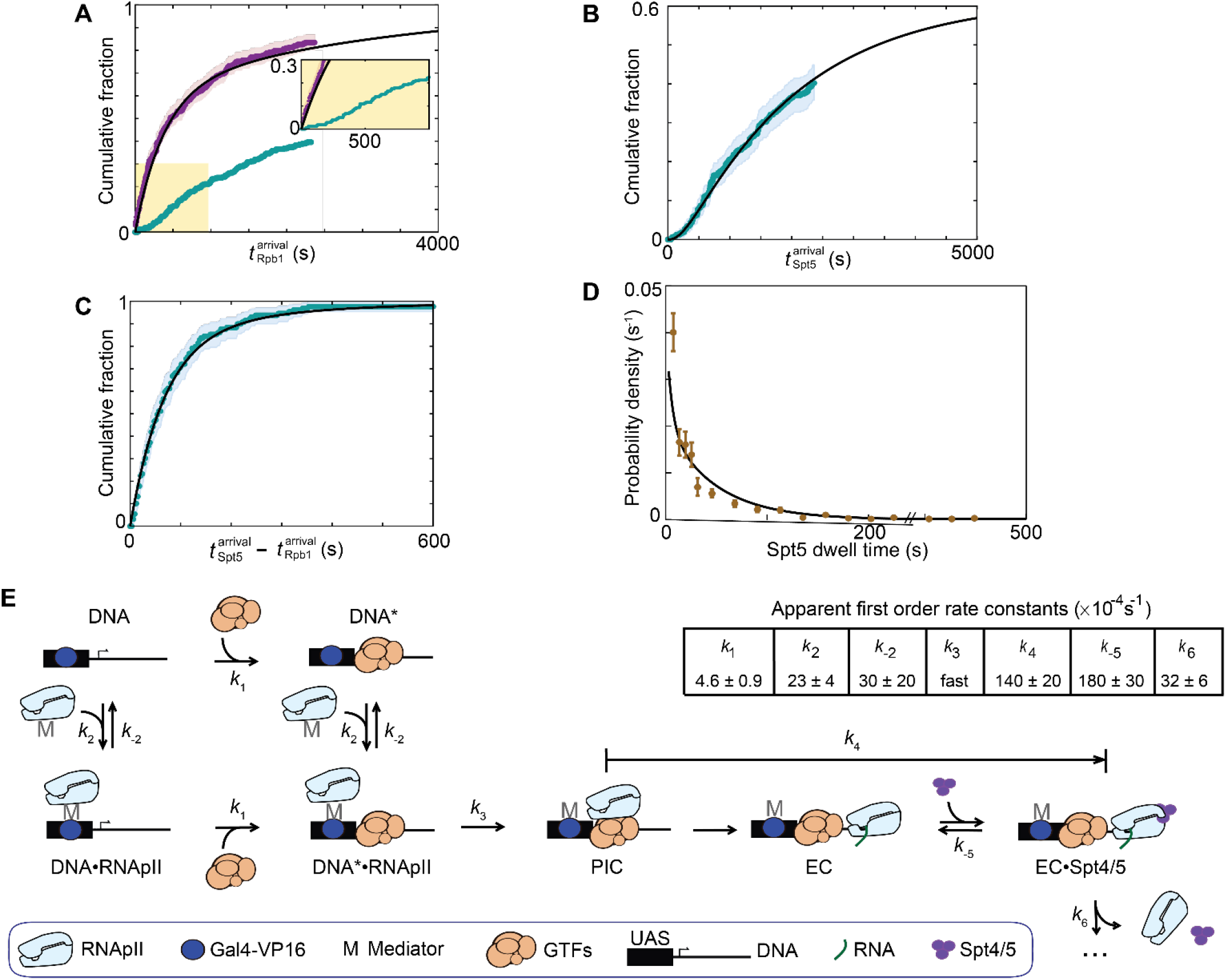
RNApII and Spt4/5 binding dynamics during activated transcription initiation in Rpb1^SNAP549^/Spt5^DHFR-Cy5^ extract. **(A)** Cumulative distribution of *N* = 268 time intervals measured from the addition of extract (*t* = 0) until the first Rpb1^SNAP549^ binding was observed at each DNA molecule. Separate curves show the distributions for all Rpb1^SNAP549^ first binding events (purple; same data as in Fig. 2C); and for the *N* = 129 first *productive* events [i.e., the first Rpb1^SNAP549^ binding event during which Spt5^DHFR-Cy5^ was also seen to bind] (turquoise). Inset: Magnified view. **(B)** Cumulative distribution of *N* = 129 time intervals from addition of extract until Spt5^DHFR-Cy5^ binding in the first productive events. **(C)** Cumulative distribution of *N* = 129 time differences between the first Spt5^DHFR-Cy5^ binding and Rpb1^SNAP549^ binding in the first productive events. Note expanded time scale compared to (A, B). Shading in A-C indicates 95% confidence interval. **(D)** Distribution (probability density ± S.E.) of Spt5^DHFR-Cy5^ dwell times during productive binding events (brown; *N* = 312). **(E)** Minimal kinetic scheme consistent with the data in (A – D), and the rate constants determined by globally fitting those data to the scheme. In the scheme shown, RNApII can bind either at the upstream Gal4 binding sites (UAS), presumed to be via an indirect interaction through Mediator and Gal4-VP16, or at the promoter, presumed to requiring a minimal set of RNApII general transcription factors (GTFs). PIC, pre-initiation complex; EC, elongation complex. Arrows represent the rate limiting step in each transformation; additional non-rate-limiting processes are not shown and are ignored in the quantitative modeling. Rate constants *k*_1_, *k*_2_, *k*_−2_, and *k*_4_ were determined by global fitting of the model to data underlying (A-C), and *k*_−5_ and *k*_6_ by fitting to the data underlying (D) and the measured partition ratio (see SI Materials and Methods). *k*_3_ was fixed to a value (1 s^−1^) faster than the experimental time resolution because the data show no evidence for a rate limitation by this step. For simplicity, this illustration does not show steps in which Rpb1^SNAP549^ or Spt5^DHFR-Cy5^ molecules bound non-specifically to the slide surface, but these kinetic processes (see **Fig. S7**) were included in the determination of the rate constants shown. Black lines in (A – D) show the distributions calculated from the model using the rate constant values in (E).

The distribution of times to first binding of Spt5^DHFR-Cy5^ while Rpb1^SNAP549^ is present, which we interpret as indicative of a full elongation complex, shows an even longer (~300 s) lag than productive Rpb1^SNAP549^ association (**Fig. 4B**, **Fig. S5**). In contrast, the time intervals between Rpb1^SNAP549^ and Spt5^DHFR-Cy5^ binding in productive events were short, with the majority less than 100 s (**Fig. 4C**). These intervals were exponentially distributed, consistent with a single rate-limiting process occurring between RNApII binding and elongation complex formation.

### Spt4/5 exchange on elongation complexes

To complete the analysis of Spt4/5 dynamics, we examined the fates of the DNA-bound RNApII•Spt4/5 complexes. The lifetimes of Spt5^DHFR-Cy5^ fluorescence spots in these productive complexes displayed an exponential distribution (**Fig. 4D**); that is, they behaved kinetically like a single molecular species. As described earlier, Rpb1^SNAP549^ and Spt5^DHFR-Cy5^ fluorescence spots sometimes disappeared from DNA simultaneously (within ±1 video frame, e.g., closed arrows in **Fig. 3A** and **Fig. S3**), consistent with dissociation of a unitary RNApII•Spt4/5 complex. However, more often Spt5^DHFR-Cy5^ fluorescence disappeared while Rpb1^SNAP549^ fluorescence remained (e.g., open arrows in **Fig. 3A** and **Fig. S3**; 77 ± 2% of 312 RNApII•Spt4/5 complexes observed). Control experiments showed that Spt5^DHFR-Cy5^ photobleaching and dissociation of the Cy5-TMP dye are not significant factors in limiting Spt5^DHFR-Cy5^ bound lifetimes (**Fig. S6**). Thus, loss of Spt5^DHFR-Cy5^ fluorescence while Rpb1^SNAP549^ remains signifies dissociation of Spt4/5 from the RNApII elongation complex. Indeed, some DNA-bound RNApII molecules that lost Spt5^DHFR-Cy5^ were subsequently seen to bind another Spt5^DHFR-Cy5^ molecule (**Figs. 3A and S3**). Of 49 DNA molecules exhibiting only a single Rpb1^SNAP549^ binding event, 15 showed multiple sequential Spt5^DHFR-Cy5^ binding and release events. Together these data indicate that Spt4/5 binding to elongation complexes is reversible, with individual elongation complexes able to undergo multiple cycles of Spt4/5 binding and release in vitro.

### Quantitative kinetic model for initiation complex formation and Spt4/5 recruitment and release

The relative kinetics of RNApII and Spt4/5 association with the DNA template (**Fig. 4A-C**) suggest a reaction mechanism in which 1) initial RNApII associations with DNA are dominated by the formation of complexes that do not go on to elongate, 2) a slower step is needed to make the promoter competent for PIC and EC formation, and 3) subsequent initiation of RNA synthesis and binding of Spt4/5 to the elongation complex are comparatively quick. To aid interpretation, we used the quantitative kinetic data to formulate a minimal scheme consistent with these results (**Fig. 4E**). The rate constants given in the figure were determined by fitting the data summarized in **Fig. 4A-D**, and the curves predicted by the model and rate constants are shown in those same panels. The data do not speak directly to the structures of the molecular species in the model, but we have proposed plausible candidates for the identities of the different DNA-associated RNApII complexes based on literature data. As illustrated (**Fig. 4E**), the scheme embodies the following assumptions:

1. Because Rpb1^SNAP549^ does not bind DNA in the control reaction lacking Gal4-VP16, we show all states of the model with this activator bound. A step encompassing Gal4-VP16 binding to the upstream Gal4 binding sites was not explicitly included since Gal4 binding is likely to be fast and occupancy high given the concentration of Gal4-VP16 used and five Gal4 binding sites in the template (39, 40).
2. The model includes an additional step (*k*_1_) needed for productive RNApII binding (i.e., the ability to eventually bind Spt4/5). This step renders the DNA competent (DNA*) for PIC and EC formation, and probably represents binding to the core promoter of a subset of the RNApII general transcription factors. In particular, it is known that at least TBP and TFIIB must be present before RNApII can proceed on the pathway to PIC formation (2, 41). Alternative simpler mechanisms lacking the non-productive RNApII binding intermediate (DNA·RNAPII) did not produce satisfactory agreement with the data.
3. We use the same rate constant for RNApII binding (*k*_2_) whether or not the minimal general transcription factors are present. The distribution of times until the first Rpb1^SNAP549^ binding to a given DNA molecule fits a single-exponential model (**Fig. 2B**), consistent with a single rate-limiting step. As RNApII recruitment is completely dependent on Gal4-VP16, we speculate that the initial binding involves association of a Mediator-RNApII complex with the DNA-bound activator. We model this step as reversible, with the dissociation rate constant (*k*_−2_) also being independent of general transcription factors.
4. While our earlier immobilized template assays suggest Gal4-VP16 promotes basal factor binding (20), the current model presumes that activator is saturating on templates. It is therefore agnostic as to whether the activator increases the rate of promoter competence (*k*_1_) in addition to increasing the rate of RNApII recruitment.
5. The *k*_1_ and *k*_2_ processes are assumed to be independent. This is equivalent to saying that the basal factor binding (or other event) that triggers promoter competence is unaffected by the presence of RNApII bound to activator at the UAS.
6. A number of possible additional steps unnecessary for fitting the data were omitted for simplicity. These include the reverse of the *k*_1_ and *k*_3_ steps (discussed below). Activator-independent binding of RNApII to general transcription factors at the promoter (DNA* → PIC in **Fig 4E**) was also omitted, given the observed strong dependence of RNApII binding on Gal4-VP16.
7. The model does not include possible binding of a second RNApII molecule at the UAS when a first RNApII molecule is already engaged as a PIC or EC. This pathway would provide a simple explanation for the occasional observation of multiple Rpb1 molecules simultaneously bound to the same DNA (**Fig. 2A**, brackets). However, when there are simultaneous multiple Rpb1^SNAP549^ bindings on a DNA template, it is impossible to conclusively link the individual binding and dissociation events. Therefore, these multiple binding events were not easily usable for kinetic modeling.

Several mechanistic insights emerge from the rate constants determined for the model. The slowest step in the mechanism is *k*_1_, the step leading to the DNA* state competent for transcription. We propose this step involves binding of one or more of the general transcription factors needed for subsequent PIC formation. Rate constant *k*_2_ is larger than *k*_1_, implying that most RNApII binding that occurs early in the reaction forms state DNA·RNApII. Because *k*_−2_ is greater than *k*_1_ by approximately six-fold, most DNA·RNApII complexes dissociate back to the DNA state, with only a fraction continuing on to the DNA*·RNApII intermediate. Thus, the model explains why most RNApII binding early in the reaction is non-productive. The two classes of RNApII binding events in the model, to DNA or DNA*, explain the difference between the exponential kinetics seen for total first Rpb1^SNAP549^ binding events (**Fig. 4A**, purple), versus the lag in the cumulative distribution of times for the first productive Rpb1^SNAP549^ binding events (**Fig. 4A**, turquoise).

Once RNApII is bound to a competent promoter (DNA*·RNApII), RNApII can in principle either dissociate to restore DNA* (*k*_−2_) or convert into a PIC (*k*_3_). The latter step may involve transfer of Mediator-RNApII from the activator to the TBP-TFIIB-TATA box complex, separation of RNApII from Mediator, or incorporation of later basal factors (TFIIF, TFIIE, and/or TFIIH). Because *k*_3_ is much greater than *k*_−2_, most complexes that get to the DNA*·RNAPII state go on to produce PICs, consistent with the idea that once all components are present, PIC formation is fast. In the model, reversibility of this step was not required to fit the data, so *k*_−3_ was set to zero for simplicity.

In our system, Spt4/5 serves to indicate the presence of an EC, but the transition from PIC to EC involves multiple molecular events, including promoter melting, release of basal factors, and binding of multiple elongation factors, all of which might precede Spt4/5 binding. Time intervals between Rpb1^SNAP549^ and Spt5^DHFR-Cy5^ bindings appear as a single exponential distribution (**Fig. 4C**), consistent with a single predominant rate-limiting step between productive RNApII binding and Spt4/5 binding. This interval is therefore modeled as a single aggregate step with rate constant *k*_4_. Future experiments may be able to better define which molecular event is rate-limiting.

## Discussion

While there has been remarkable recent progress in our understanding of the structures of RNApII transcription complexes, far less is known about the dynamics of transitions between the different structural states. Here we report the use of CoSMoS to directly observe the dynamics with which single fluorescently labeled molecules of RNApII, the elongation factor Spt4/5, and the mRNA binding protein Hek2 interact on a transcription template DNA. In contrast to most other single molecule studies, these experiments were carried out in nuclear extracts, which should more closely approximate conditions in the nucleus than do systems reconstituted from RNApII and a subset of purified transcription factors. Using a template carrying multiple Gal4 binding sites upstream of the well-characterized *CYC1* core promoter, we observe that RNApII binding and subsequent transcription are strongly dependent on the activator Gal4-VP16. Analysis of RNApII binding dynamics reveals distinct classes of productive and non-productive binding events and suggests there is a rate limiting process that makes the promoter competent for PIC formation. Kinetic modeling suggests that initial binding of RNApII to the activator, possibly as an RNApII-Mediator complex, is relatively fast. The slower competence step is likely to reflect association of one or more general transcription factors. Future CoSMoS experiments can rigorously test these modeling-derived hypotheses by labeling additional components of the transcription machinery.

The reported experiments capture the transition from initiation to elongation, which we previously analyzed by quantitative mass spectrometry (20). To detect EC formation in the single-molecule experiments, we initially tried hybridization of fluorescently labeled probe oligonucleotides to the nascent RNA, as has been done single-molecule transcription experiments using purified factors (27, 42, 43). Probe detection of nascent transcript was ineffective in extract, perhaps because RNA-binding proteins interfere with hybridization. As an alternative, we fluorescently labeled the endogenous mRNA binding protein Hek2, which is highly enriched in ECs on this template (20). We observed that Spt4/5 occupancy is strongly correlated with Hek2 presence (**Fig. 2E**), and is nucleotide dependent (**Fig. 3B**), and therefore conclude that Spt4/5 binding is confined to ECs, consistent with our mass spectrometry observations.

The kinetics of RNApII basal transcription initiation (i.e., initiation that occurs in the absence of activators) has been previously examined in vitro, including by single-molecule methods (41, 42, 44–46). Such studies typically observe transcription from only a small fraction of template molecules, although this can be greatly improved by pre-formation of PICs before initiating RNA synthesis (47–49). Here we focused on the dynamics of activated transcription, which is thought to be the dominant mode of gene expression in vivo. As observed for Gal4-dependent promoters in vivo (50), transcription in our system is highly activator-dependent. High efficiency is suggested by the observation that in most experiments which included Gal4-VP16 the majority of template molecules bound RNApII.

Chromatin immunoprecipitation experiments show Spt4/5 crosslinking throughout transcribed regions (4, 51). This result is consistent with either stable incorporation of Spt4/5 into the EC or a dynamic interaction in which Spt4/5 dissociates and reassociates during synthesis of a single transcript. In the CoSMoS experiments, when we saw Spt4/5 dissociate before RNApII it was not unusual to observe re-binding of another Spt4/5 molecule to RNApII, likely indicating dynamic interaction. The average length of a yeast pre-mRNA is 1.5 kb (52), so 45 to 90 s are needed for RNApII to transcribe an average gene if the mean elongation rate is roughly 1 to 2 kb min^−1^ (53, 54). This time range aligns with the average time measured for Spt5^DHFR-Cy5^ to dissociate from elongation complexes, 1 / *k*_−5_ = 56 ± 9 s (**Fig. 4E**). We speculate that Spt4/5 binding times may be tuned to match the average EC lifetime. Thus, a single Spt4/5 molecule would stay attached to the EC throughout a normal, unimpeded transcription cycle. On the other hand, an excessively long interval of pausing or backtracking may cause the EC to persist beyond the average lifetime of Spt4/5 binding, allowing dissociation of Spt4/5 and favoring termination of these RNApII molecules.

Transcription activators, Mediator, and RNApII are reported to participate in self-associated clusters or condensates in mammalian cells, and it is speculated that such association may play an important role in transcription activation (ref. (55) and refs. therein). In our experiments we see no evidence for clusters or condensates containing multiple RNApII molecules. Instead, we observe association of individual RNApII molecules with DNA, followed by the progression of these molecules to active ECs. Our results do not exclude the involvement of condensates in vivo, where nuclear protein concentrations are substantially higher than those in our extracts. Rather, they demonstrate that polymerase condensate formation is not required for activator stimulation of RNApII transcription, even in a strongly activator-dependent system like the one studied here.

In recent years, researchers have reconstituted numerous molecular complexes representing intermediates in RNApII transcription initiation and transcript elongation, processing and termination. High resolution three-dimensional structures of the complexes show how transcription factors, DNA template and RNApII interact, and the structures suggest mechanisms by which the interactions are remodeled to coordinate and regulate mRNA production. In contrast to this wealth of structural information, we know less about the arrangement of these intermediates in the reaction pathway and even less about the dynamics of the reactions that drive the transcription machinery through successive intermediates. The approach used here represents a generally applicable tool to elucidate the dynamic mechanisms of transcript production by RNApII and its regulation.

## Materials and Methods

### Yeast strains, plasmids and oligonucleotides

*S. cerevisiae* strains used in this study are listed in **Table S5**. SNAP- or DHFR-tagged fusion strains were constructed by PCR amplification of a cassette encoding the SNAP/DHFR protein and a selectable marker. The DHFR-containing plasmid was generously provided by Aaron Hoskins (22). Oligonucleotides (IDT Ultramers) for PCR had 50 nt of homology to the target gene C-terminal coding region and 20 nt of homology to the fusion cassette. The amplified fragment was transformed into yeast with selection for the marker. In-frame fusion protein expression and stability was confirmed by immunoblotting for the target protein (e.g., **Fig. S1A, B**), and in the case of SNAP fusions, by SDS-PAGE after labeling with SNAP-Surface 549 (New England Biolabs). The strains had doubling times similar to their parental strains YF702 or YF4 (**Fig. S1E**).

### Yeast nuclear extracts

Yeast nuclear extracts were prepared as previously described (17). For SNAP fusion strains, the protocol was modified in that nuclear protein pellets were resuspended in 1–2 mL of buffer C′ (20 mM HEPES, pH 7.6, 10 mM MgSO_4_, 1 mM EGTA, 10% glycerol, 3 mM DTT, and 1 μg ml^−1^ each of aprotinin, leupeptin, pepstatin A, and antipain). (In pilot experiments, the 20% glycerol concentration and PMSF used in (17) were found to reduce SNAP labeling.) To fluorescently label extracts with SNAP fusion proteins, SNAP-Surface 549 was added with gentle vortexing to the resuspended pellet at a final concentration of 0.5 μM (unless otherwise specified), and the mixture was incubated at 4°C for 1 hr on a rotator. All extracts were dialyzed against 3 × 0.5 L buffer C′ supplemented with 75 mM (NH_4_)_2_SO_4_. For SNAP-labeled extracts, unreacted SNAP-Surface 549 was further depleted as described (56) with some modifications. Specifically, 0.25 volumes of SNAP-agarose beads (see SI Methods) was added to the dialyzed extract and incubated at 4°C for 1 hr on a rotator, after which the beads were removed by centrifugation at 1,000 × *g* for 2 min at 4°C. All extracts were aliquoted, frozen in liquid N_2_, and stored at –80°C. Labeling of SNAP fusion proteins and depletion of free dye was confirmed by SDS-PAGE followed by fluorescence imaging on a Typhoon imager (GE Healthcare). Extracts were checked to verify transcription activity (e.g., **Fig. S1C, D**). If DNA-specific Spt5^DHFR-Cy5^ binding occurred at a frequency <0.5 × 10^−4^ s^−1^, the extract was considered to have inactive or untagged Spt5 and was not used.

### DNA template

Transcription template (**Fig. 1A**) was prepared by PCR from plasmid SB649 (20) using primers 5′-biotin-GTTGGGTAACGCCAGGG-3′ and 5′-Alexa488-GGAAACAGCTATGACATG-3′ (IDT) and Platinum Taq DNA Polymerase (Invitrogen). The PCR product was purified using DNA SizeSelector-I SPRI magnetic beads (Aline Biosciences) according to the manufacturer’s instructions. The template contained five tandem repeats of 5′-CGGAGGACAGTACTCCG-3′, a consensus Gal4 binding sequence (57).

### CoSMoS transcription experiments

Single-molecule fluorescence microscopy experiments were conducted at 20-23° C, 1 frame s^−1^ (except as otherwise noted) on a micromirror total internal fluorescence microscope at excitation wavelengths of 488, 532, and 633 nm (58) using laser powers incident to the micromirror of 700, 400, and 200 μW respectively. Focus was automatically maintained as described (59). Transcription reactions were observed in glass flow chambers passivated with a mPEG-SG2000:biotin-PEG-SVA5000 (Laysan Bio) 200:1 w/w mixture as described in (27). Streptavidin-coated fluorescent beads (T-10711, Molecular Probes) to serve as fiducial markers for stage drift correction (59) were added to the flow chamber at a dilution of ~1:400,000 in transcription buffer (20 mM potassium acetate, 20 mM HEPES, pH 7.6, 1 mM EDTA, 5 mM magnesium acetate) supplemented with 1 mg mL^−1^ bovine serum albumin (TB+BSA). The chamber was subsequently incubated with 0.013 mg/mL NeutrAvidin (31000, Thermo Fisher) in TB+BSA for 45 s and then flushed with three chamber volumes (3 × 20 μL) of TB+BSA. We then introduced ~10 pM DNA^488^ in TB+BSA and incubated until 200-400 fluorescent spots were visible under 488 nm excitation. After moving the stage to a new field of view, the transcription reaction was started by introducing a mixture of yeast nuclear extract, TB, and other reagents with the following final concentrations: extract (6.6 mg mL^−1^ final protein concentration), O_2_-scavenging system (60) (0.9 units mL^−1^ protocatechuate dioxygenase, 5 mM protocatechuate), 2 mM DTT, triplet state quenchers (61) (0.5 mM propyl gallate, 1 mM Trolox, 1 mM 4-nitrobenzyl alcohol, 0.5% DMSO), ATP regeneration system (10 mM phosphocreatine. 0.1 mg mL^−1^ [35 units ml^−1^] creatine kinase), 20 nM Cy5-TMP (22) (only if Spt5^DHFR^ was present), zero or 4 μg mL^−1^ Gal4-VP16 activator, nucleotides (400 μM each ATP, CTP and UTP plus 100 μM 2’-O-methyl-GTP to aid in stalling the RNApII at the end of the G-less cassette), and buffer components (100.5 mM potassium acetate, 24.1 mM HEPES, pH 7.6, 0.4 mM Tris-acetate, 1 mM EDTA, 5 mM magnesium acetate, 2 mM MgSO_4_, 0.2 mM EGTA, 15 mM (NH_4_)_2_SO_4_, 4.25% glycerol). We monitored labeled extract component association to and dissociation from the recorded surface-tethered DNA^488^ locations by exciting individually (one-color experiments) or simultaneously (two-color experiments) with the 532 and 633 nm lasers.

For the –NTPs control experiments, nucleotides and ATP regeneration components were omitted and residual NTPs were depleted by the addition of 20 mM glucose and 40 units mL^−1^ hexokinase (Sigma H4502). In inhibitor controls, α-amanitin was added at 10 mg mL^−1^. For the photobleaching control experiment (**Fig. S6E**), we conducted three separate transcription experiments with different duty ratios of the 633 nm laser at 200 μW.

### Data analysis and kinetic modeling

See SI Materials and Methods.

## Acknowledgements

We thank Judy Ciyue Shen and Yujin Chun for constructing yeast strains used in this work, and members of the Buratowski and Gelles labs for advice and comments on the manuscript. Research reported in this publication was supported by the National Institute of General Medical Sciences and the National Cancer Institute of the National Institutes of Health (NIH) under award numbers R01GM056663 (to S.B.) R01GM081648 (to J.G.) and R01CA246500 (to S.B. and J.G.). The content is solely the responsibility of the authors and does not necessarily represent the official views of the National Institutes of Health.

## SI Materials and Methods

### CoSMoS Data Analysis

Image analysis was done using custom software and algorithms for automatic spot detection, spatial drift correction, and colocalization as described (59). All reported statistical uncertainties are standard errors unless otherwise indicated.

To analyze RNApII association to template DNA locations (**Fig 1C, D; Table S1**), we used a kinetic model that assumed a time-independent apparent first-order rate constants *k*_on_ for specific association of RNApII with the DNA template (including any bound proteins) and *k*_p,n_ for non-specific association of RNApII to the chamber surface. As previously (59), we further assumed that only an active fraction *A*_f_ of DNA^488^ surface-bound molecules were capable of RNApII binding. As a preliminary step in the analysis of and experiment, we measured the time to first observed binding of labeled RNApII to randomly selected no-DNA locations and performed maximum likelihood fitting using an exponential model to determine *k*_p,n_. We then fit the times of first binding to DNA^488^ locations using a model that included both exponential non-specific binding (with *k*_p,n_ held fixed at the previously determined value) and exponential specific binding (at rate *k*_on_ to fraction of DNA^488^ molecules *A*_f_). Standard errors of fit parameters were determined by bootstrapping. Analysis was restricted to first binding events for the reasons described previously (59).

Protein binding frequencies were calculated as defined in (59). To measure DNA-specific binding frequencies of Spt5 or Hek2 (**Fig. 3B, D; Tables S2 and S3**), we subtracted the total frequency of Spt5^DHFR-Cy5^ or Hek2^SNAP549^ binding to no-DNA locations from the total frequency of Spt5^DHFR-Cy5^ or Hek2^SNAP549^ binding to DNA locations.

DNA-specific fractional occupancy of DNA molecules by Spt5^DHFR-Cy5^ with or without a colocalized Hek2^SNAP549^ molecule (**Fig. 3E**; **Table S4**) was calculated as in (23). Standard errors of the fractional occupancies were calculated by delete-1 jackknife resampling: briefly, the fractional occupancy was recalculated for the dataset *n* times, each time eliminating a single observation, for data containing *n* instances of Spt5^DHFR-Cy5^ colocalized on DNA molecules with or without a simultaneously colocalized Hek2^SNAP549^ molecule. The standard error of the initially measured fractional occupancy was taken to be the standard deviation of the fractional occupancies calculated with the resampled data sets.

### Global kinetic modeling

We formulated a minimal kinetic model (shown in summary form in **Fig. 4E** and in full detail in **Fig. S7**) to account for the observed single molecule dynamics. The model includes both specific protein-protein and protein -nucleic acid interactions as well as non-specific binding of Spt5^DHFR-Cy5^ and Rpb1^SNAP549^ to the chamber surface. The non-specific rate constants were determined by measuring times to first appearance of the proteins at randomly selected no-DNA locations and fitting these data to exponential distributions (59). These non-specific binding rate constants *k*_p,n_ and *k*_s,n_ were then held fixed in the subsequent analysis. To determine parameters *k*_1_, *k*_2_, *k*_−2_, *k*_4_, and the active fraction *A*_f_, we used maximum likelihood methods to globally the fit the kinetic model to three sets of measurements whose distributions are plotted in **Fig. 4A** (purple), **Fig. 4B** and **Fig. 4C**. Numerical solutions to the kinetic scheme (**Fig. S7**) provided probability density functions (pdfs) for these three sets of binding data, and those numerical pdfs were used for the maximum likelihood optimization of the rate constants and *A*_f_.

To derive the likelihood function, we treated each individual reaction in **Fig. S7** as first order or pseudo-first order and solved the set of first-order linear differential equations corresponding to the complete model (**Fig. S7**) using numerical methods (62) implemented in custom Matlab programs. For a given a set of parameter values (*k*_1_, *k*_2_, *k*_−2_, *k*_4_, and *A*_f_) and initial conditions *f*_**species**_(*t* = 0) (i.e., the fraction of template DNA molecules in each species in **Fig. S7** at time *t* = 0), the numerical solution yielded fractions *f*_**species**_(*t*) for each species for all *t* > 0. These were then used to calculate pdfs *p*(*t*) for each type of experimental observation as follows:

To determine *p*_1_(*t*) for the *n*_1_ observed times *t*_i_ to first binding of Rpb1^SNAP549^ (**Fig. 4A**, purple), we set initial conditions *f*_DNA_(0) = *A*_f_ and *f*_I_(0) = (1 – *A*_f_), fixed *k*_−2_ = *k*_3_ = *k*_s,n_ = 0, and defined

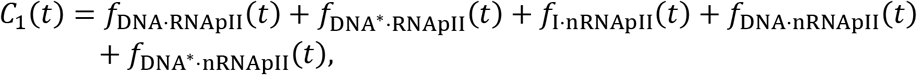

and

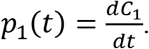

The contribution of these data to the overall log-likelihood is then

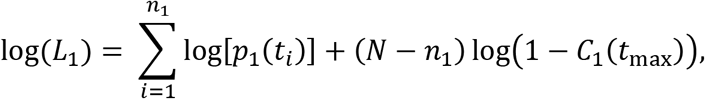

where *N* is the number of DNA^488^ locations, *n*_1_ is the number of DNA sites at which we observe a first binding of RNApII, and *t*_max_ is the duration of the experimental record. The second term in the above equation accounts for locations at which no RNApII binding was observed.

To determine *p*_2_(*t*) for the *n*_2_ observed times *t*_i_ to first binding of an Spt5^DHFR-Cy5^ molecule while Rpb1^SNAP549^ is present (**Fig. 4B**), we set *f*_DNA_(0) = *A*_f_, *f*_I_(0) = (1 – *A*_f_), *k*_−5_ = *k*_6_ = *k*_p,n_ = 0 and defined

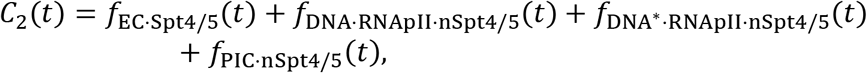

and

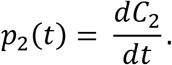

The contribution of these data to the overall log-likelihood is then

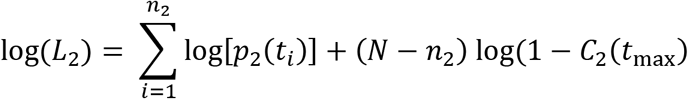

To determine *p*_3_(*t*) for the *n*_3_ observed time differences *t*_i_ between Rpb1^SNAP549^ binding in the first productive events and the first Spt5^DHFR-Cy5^ binding in those events (**Fig. 4C**), we set *f*_DNA_(0) = *A*_f_, *f*_I_(0) = (1 – *A*_f_), *k*_−5_ = *k*_6_ =*k*_p,n_ = 0, and defined

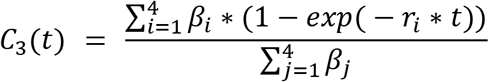

where

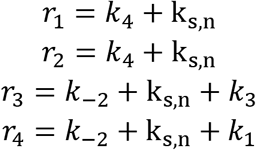

and the β_*i*_ are the asymptotic fractional occupancies

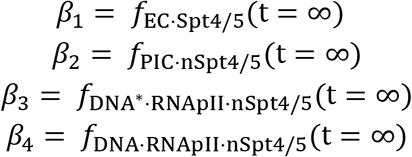

determined from a run of the numerical model to long times, and

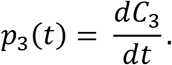

The contribution of these data to the overall log-likelihood is then

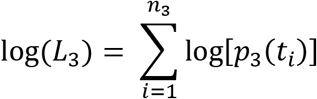

Finally, the overall log-likelihood is computed as

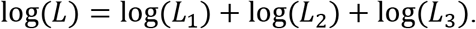

In preliminary analyses, we found that the optimized value of *k*_3_ was always faster than the time resolution of the experiment, so for simplicity its value was held fixed in the final analysis at *k*_3_ = 1 s^−1^ without diminishing the agreement of model and data. Similarly, preliminary fits gave negligible rates for the reverse of the *k*_3_ step, so this step was taken to be irreversible in the final analysis. Without the *k*_3_ step, the fit for the time to first Spt5^DHFR-Cy5^ association during Rpb1^SNAP549^ association failed to capture the initial lag present in the data.

To obtain *k*_−5_, and *k*_6_, we first fit the Spt5^DHFR-Cy5^ lifetimes that occur during Rpb1^SNAP549^ association (**Fig. 4D**) with a maximum likelihood single-exponential model that takes into account nonspecific surface binding (59), yielding the aggregate dissociation rate constant *k_app_* = *k*_−5_ + *k*_6_. The lifetimes include events in which RNApII and Spt4/5 molecules departed the DNA simultaneously as a complex, as well as those in which the Spt4/5 molecule dissociated prior to RNApII dissociation. The value from fitting these pooled data (*k_app_* = 0.021 ± 0.003 s^−1^) was identical within experimental uncertainty to the values obtained by fitting only prior (*k*_app,prior_ = 0.021 ± 0.003 s^−1^) or only simultaneous (*k*_app,simul_ = 0.018 ± 0.003 s^−1^) dissociation events, consistent with the proposal that both types of events arise from a single species (EC•Spt4/5 in **Fig. 4E**). We measured the fraction of Spt5^DHFR-Cy5^ molecules that dissociated before the accompanying Rpb1^SNAP549^ molecule as *g*_prior_ = 0.85 ± 0.02 (*N* = 312). We then calculated the rate constants as

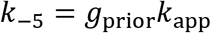

and

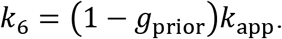

### DHFR•Cy5-TMP stability/photostability in yeast nuclear extract

Plasmid eDHFRL28C, encoding of His_6_-tagged *E. coli* DHFR (L28C mutant) (33), was constructed by Q5 site-directed mutagenesis (New England Biolabs; primers 5′-GCCTGCCGATTGCGCCTGGTTTA-3′ and 5′-AGGTTCCACGGCATGGCGTTTTC-3′) of the corresponding wild-type construct, which was generously provided by L. Hedstrom (63). The His_6_-DHFR(L28C) protein was expressed in BL21(DE3) pLysS cells grown to mid-log phase, induced with 1 mM IPTG, and grown for 3 hr at 37 °C. Cell pellets were resuspended in lysis buffer (0.5 M NaCl, 20 mM Na_2_HPO_4_, and 10 mM imidazole, pH 8.0) and sonicated on ice. The resulting lysate was clarified (70,000 × *g* for 30 min at 4 °C) and the supernatant was loaded onto a HisTrap nickel-NTA column and eluted with a 10 - 400 mM imidazole gradient in lysis buffer. Purified protein was dialyzed into PBS (137 mM NaCl, 2.7 mM KCl, 10 mM Na_2_HPO_4_, 1.8 mM KH_2_PO_4_, pH 7.4), snap frozen in liquid nitrogen, and stored at −80 °C.

To observe the lifetimes of DHFR^Cy5-TMP^ fluorescence spots, that is, the time intervals that terminated by dissociation of the non-covalent complex, fluorescence blinking, or photobleaching, we first preincubated a neutravidin-coated flow chamber with 10 nM anti-penta-His antibody (Qiagen), 10 nM His_6_-DHFR(L28C), and 20 nM Cy5-TMP in TB+BSA. After accumulation of a sufficient surface density of fluorescent spots, the chamber was washed twice or three times with TB+BSA. To initiate dissociation of Cy5-TMP from the surface-tethered protein, we added yeast nuclear extract (without any DHFR-tagged proteins) in a solution of the same composition used in the CoSMoS transcription experiments but supplemented with 2 μM unlabeled TMP competitor to block rebinding of any Cy5-TMP released into solution. Lifetimes of DHFR^Cy5-TMP^ spots present at the beginning of the recording were measured as the time of the first image with no spot at that location. In rare cases (69 of 797 spots observed over two experimental replicates), a spot displayed more than one fluorescence intensity level over the duration of the 25 min recording, presumably because of multiple DHFR^Cy5-TMP^ complexes initially present at the same location. These locations were excluded from further analysis.

### Bulk *in vitro* transcription assay

*In vitro* transcription was performed as described previously (17) with some modifications, using SB649 plasmid DNA (20) as the template. After a 5 min preincubation of 6.7 ng/μL (0.3 μM) of Gal4-VP16 with the template DNA, nuclear extract and 400 μM each of ATP, CTP, UTP along with [α-^32^P] UTP were added to the transcription mixture and incubated for 45 min at room temperature. Transcripts were recovered after treatment with RNaseT1 and resolved on an 8 M urea 5% polyacrylamide gel.

### Yeast growth assays

Overnight cultures from single colonies were diluted five-fold and then grown for three hours in YPD. After the growth, cultures were diluted with YPD to OD_600_ = 0.002. Five 200 μL wells of each strain were grown with agitation at 30 °C for 80 hours in a Tecan microtiter plate reader. Growth rates was determined by fits to the linear portion of the log(OD_600_) vs. time curves.

### SNAP-agarose beads

Agarose beads conjugated with His-tagged SNAPf protein (64) (His-SNAPf) were prepared as described (56) with some modifications. Specifically, BL21(DE3) cells containing pET24b-6His-fSNAP (56, 65) were grown at 37°C in LB medium containing 50 μg/mL kanamycin to an OD_600_ of 0.6–0.8. Protein expression was induced with 0.4 mM IPTG. Cells were further cultured at 18°C overnight and harvested by centrifugation at 5,000 rpm for 20 min at 4°C. Pellets were resuspended in lysis buffer (20 mM HEPES, pH 7.6, 500 mM NaCl, 5 mM 2-mercaptolethanol, 10% glycerol, 1 μg ml^−1^ each of aprotinin, leupeptin, pepstatin A, and antipain) and lysed by sonication. Soluble extracts were collected and incubated with pre-washed Ni^2+^-NTA-agarose resin (Gold Bio) for 1 hr on a rotator at 4°C. Bound proteins were eluted with elution buffer (lysis buffer supplemented with 300 mM imidazole). Eluted protein was dialyzed against PBS (137 mM NaCl, 2.7 mM KCl, 10 mM Na_2_HPO_4_, and 1.8 mM KH_2_PO_4_, pH 7.4, with 10% glycerol and 3 mM DTT, yielding protein at 15–20 mg/mL. NHS-activated agarose resin (Thermo Scientific) was incubated with purified His-SNAPf protein at a ratio of 90 mg resin to 10 mg protein for 2–3 hr at room temperature, quenched with 1 M Tris-HCl, pH 7.5 for 30 min at room temperature, then then washed with 10 mL PBS using a Pierce centrifuge column (Thermo Fisher Scientific, 89897). Prior to use the beads were equilibrated with buffer C′ supplemented with 75 mM ammonium sulfate.

## Supplementary Figures

**Figure S1:**
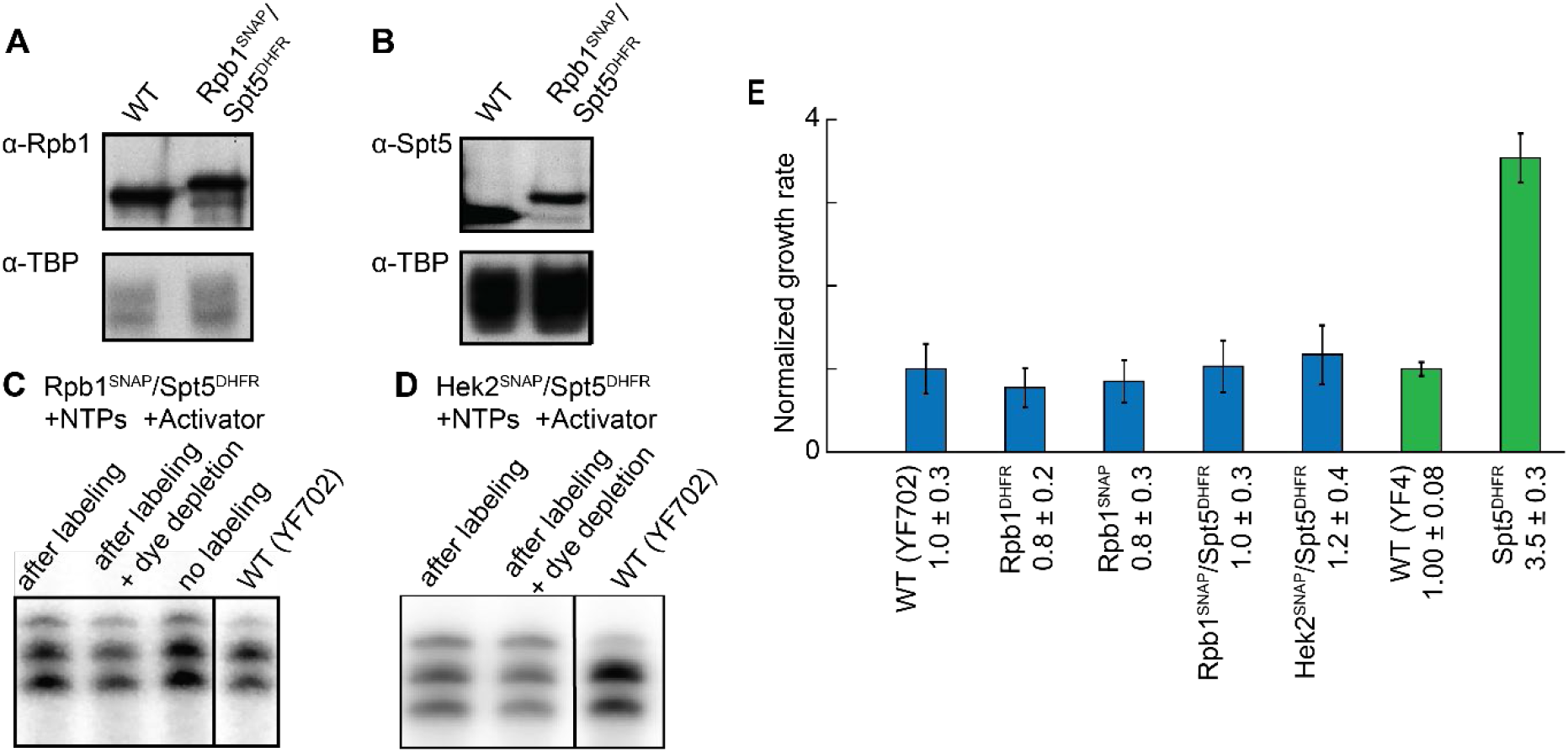
Tagged yeast strains have growth rates, nuclear extract protein expression levels, and transcription activities similar to wild-type. **(A)** Western blot showing that the concentrations of Rpb1^SNAP^ in Rpb1^SNAP^/Spt5^DHFR^ strain (YSB3403; see **Table S5**) (lane 2) and Rpb1 in parental strain (WT; YF702) are similar. TATA binding protein (TBP) was used as a loading control. Rpb1 was detected by monoclonal antibody 8WG16 (66) and TBP by a polyclonal anti-TBP antibody (67). **(B)** Western blot showing that the concentration of Spt5^DHFR^ in Rpb1^SNAP^/Spt5^DHFR^ strain is similar to that of Spt5 in the parental strain. The anti-Spt5 antibody (68) was provided by Grant Hartzog. **(C and D)** RNA produced in bulk from plasmid transcription by Rpb1^SNAP^/Spt5^DHFR^ (C) and Hek2^SNAP^/Spt5^DHFR^ (D) nuclear extracts in comparison with extracts from corresponding parental strains (WT; see **Table S5**). Where indicated, extracts were labeled with 0.5 μM SNAP-Surface 549 and subsequently depleted of unreacted dye. Bulk *in vitro* transcription reactions were performed with 400 μM each ATP, CTP, and UTP supplemented with [α-^32^P]- UTP, separated by urea polyacrylamide gel electrophoresis, and visualized by autoradiography. The lower two bands in each lane are the transcripts from the two major transcription start sites of the template. The same volume of nuclear extract preparation was used in each reaction (10 μL in C and 8 μL in D); all extract preparations were 30-40 mg/mL protein. **(E)** Growth rates (mean ± S.E.; *N* = 5) of tagged strains relative to the corresponding parental (WT) strain. Colors denote correspondence between parental (YF702, blue; YF4, green) and tagged strains.

**Figure S2:**
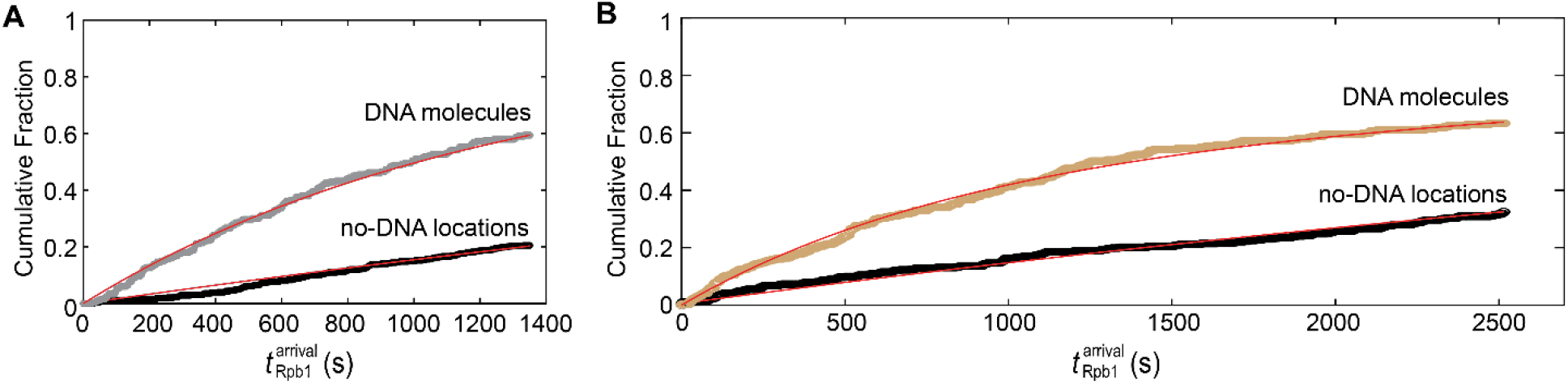
Rpb1 binding to DNA in control experiments. Graphs show cumulative distributions of time intervals prior to the first Rpb1^SNAP549^ association seen at each DNA^488^ molecule (gray/brown) and at no-DNA^488^ control locations (black), along with fits (lines) to an exponential binding model. Fit parameters are given in **Table S1**. Conditions: **(A)** +NTPs, +Gal4-VP16, 10 μg μL^−1^ α-amanitin, Rpb1^SNAP549^ extract; or **(B)** – NTPs, +Gal4-VP16, Rpb1^SNAP549^/Spt5^DHFR-Cy5^ extract.

**Figure S3:**
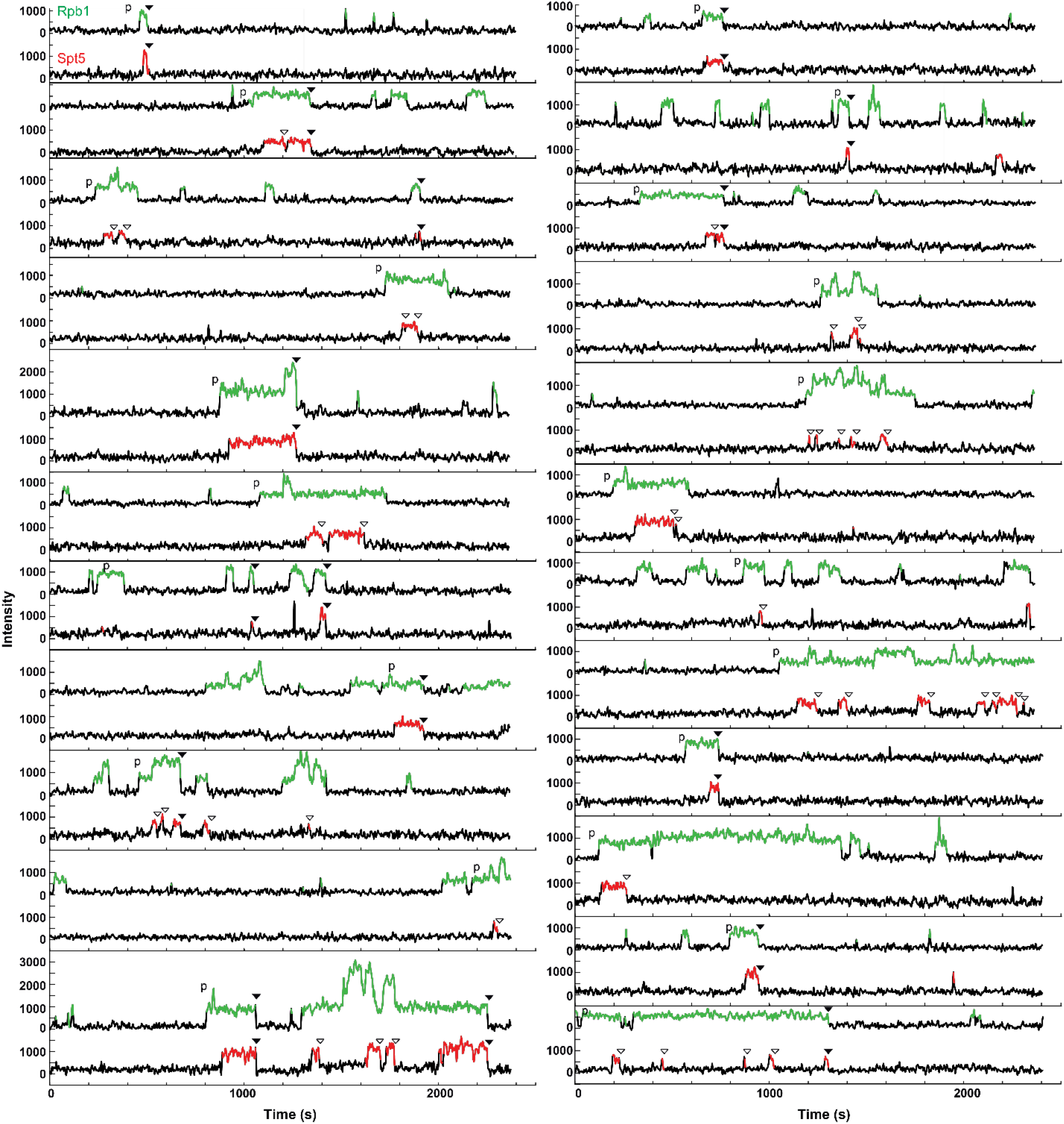
Additional example time records of Rpb1^SNAP549^ and Spt5^DHFR-Cy5^ fluorescence at 23 individual DNA^488^ locations. These records are taken from the Rpb1^SNAP549^/Spt5^DHFR-Cy5^ experiment shown in **Fig. 1.** Records are plotted as in **Fig. 3A**. The first Rpb1^SNAP549^ binding event on each DNA molecule that was designated productive (in that a Spt5^DHFR-Cy5^ molecule was observed while Rpb1^SNAP549^ was present) is marked (‘p’). Each Spt5^DHFR-Cy5^ departure is marked according to whether it occurred simultaneously with (closed triangles) Rpb1^SNAP549^ departure or not (open triangles).

**Figure S4:**
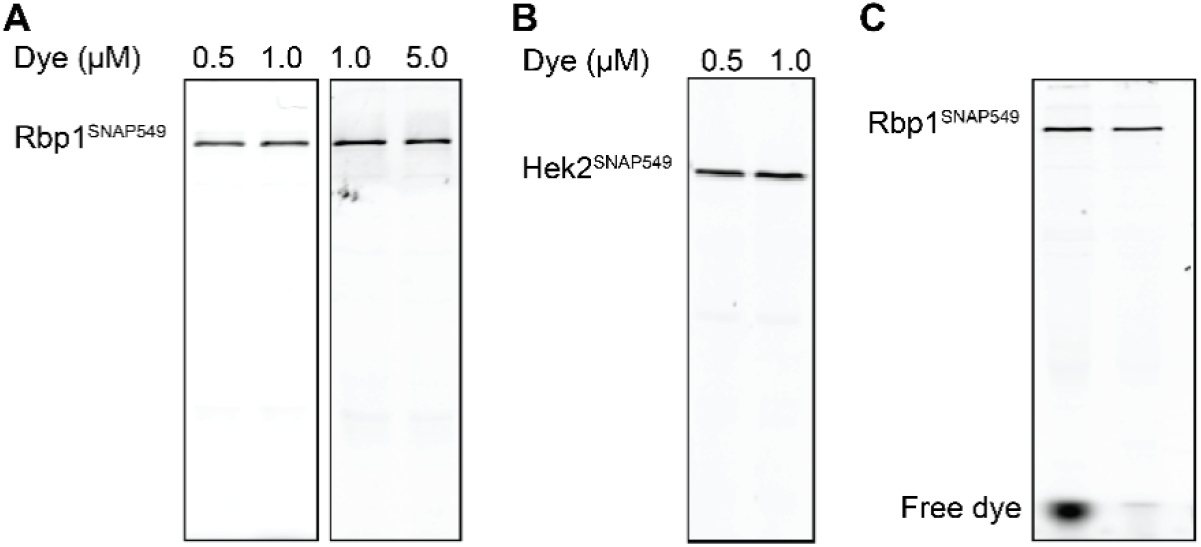
Labeling efficiency of Rpb1^SNAP549^ in nuclear extracts. Panels show SDS PAGE gels scanned for SNAP-Surface 549 fluorescence intensity. **(A)** Rpb1^SNAP549^/Spt5^DHFR^ yeast nuclear extract prepared with different concentrations of SNAP-Surface 549 dye and depleted of residual dye as described (see Materials and Methods). A concentration of 0.5 μM SNAP-Surface 549 was judged sufficient for maximal labeling. **(B)** Same as (A) but for Hek2^SNAP549^/Spt5^DHFR^ extract. Again, a concentration of 0.5 μM SNAP-Surface 549 was judged sufficient for maximal labeling. **(C)** Effectiveness of the dye depletion protocol at removing unincorporated dye after extract labeling. Lanes contain Rpb1^SNAP^/ Spt5^DHFR^ extract labeled with 0.5 μM SNAP-Surface 549 before (left) and after (right) dye depletion.

**Figure S5:**
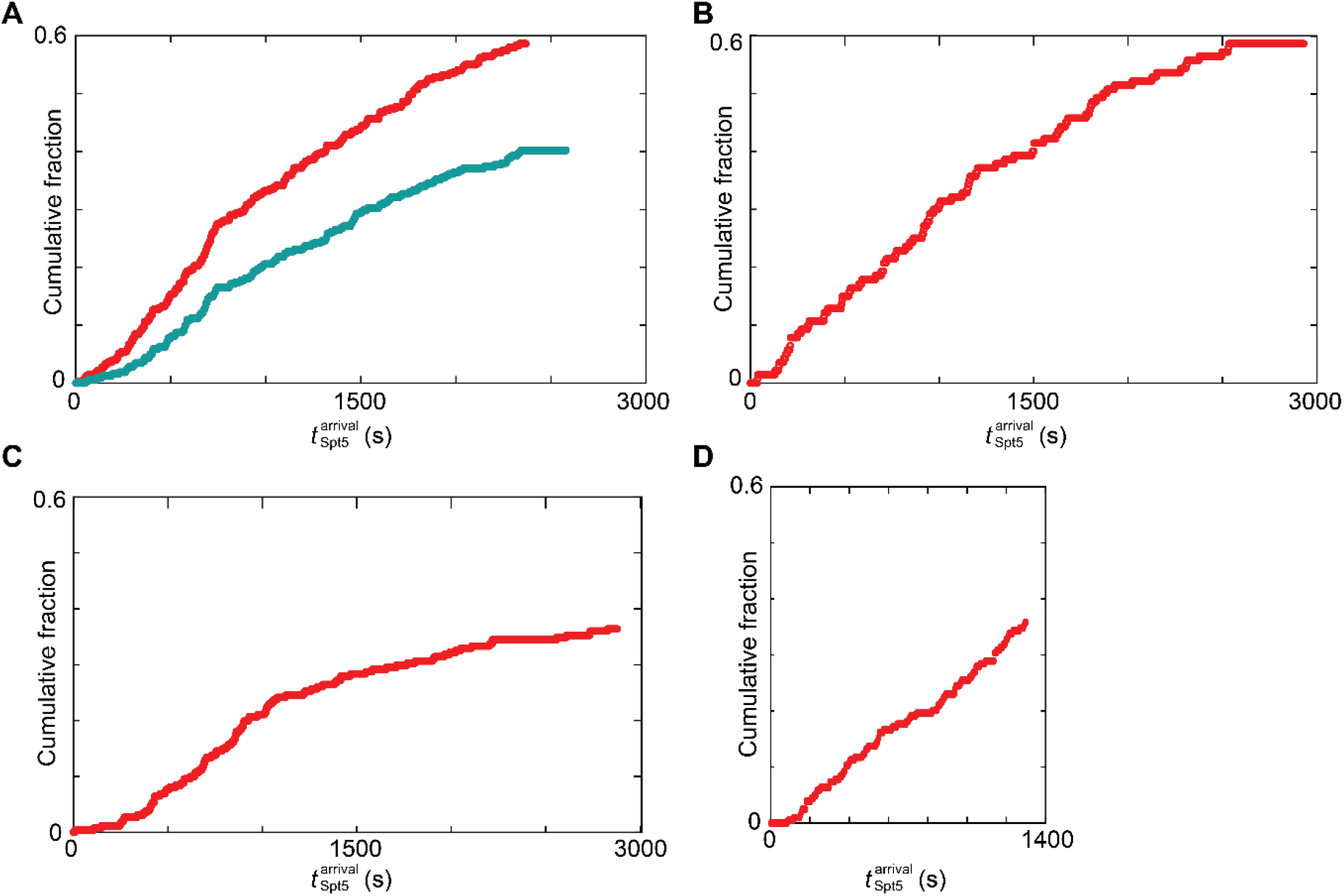
Cumulative distributions of time intervals from addition of extract until to first Spt5^DHFR-Cy5^ binding on each DNA observed in four different experiments. Data are shown either for the entire set of first Spt5 ^DHFR-Cy5^ appearances in the experiment (red) or for only Spt5 ^DHFR-Cy5^ appearances that occur during the first productive Rpb1^SNAP549^ binding event (turquoise; same data as in **Fig. 4B**). Extracts were Rpb1^SNAP549^/Spt5^DHFR-Cy5^ **(A)** or Spt5^DHFR-Cy5^ **(B-D)**. All distributions display a lag, but there are some quantitative differences in Spt5^DHFR-Cy5^ kinetics between experimental replicates (**Table S2**).

**Figure S6:**
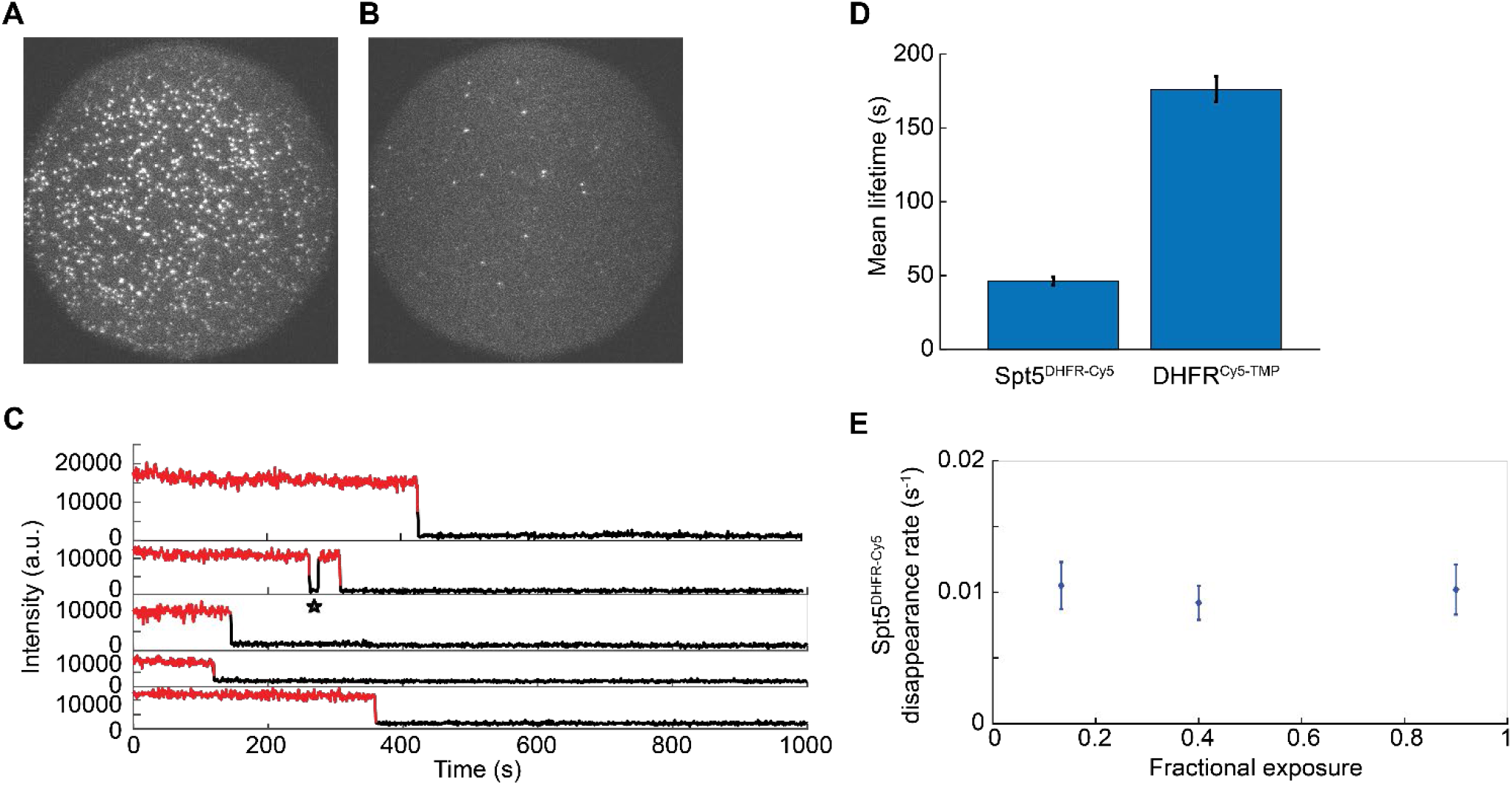
Stability and photostability of DHFR^Cy5-TMP^ and Spt5^DHFR-Cy5^ in yeast nuclear extract. **(A)** Emission of His_6_-DHFR(L28C) immobilized on an anti-penta-His antibody-coated slide surface that were preincubated with Cy5-TMP. A total of 535 fluorescent spots are visible in this 65 × 65 μm field of view immediately after adding a yeast extract without any DHFR-tagged proteins in solution conditions identical to those used in the CoSMoS transcription experiments but with the addition of 2 μM unlabeled TMP competitor. **(B)** Negative control. Same as (A), but no DHFR protein was added; only 21 non-specifically bound Cy5-TMP molecules are visible. **(C)** Example intensity records of individual DHFR^Cy5-TMP^ spots, from the experiment shown in (A). Most spots disappeared irreversibly; only 9% (68 of 728 spots from two replicate experiments) of spots showed a candidate blinking event (asterisk) in which a spot disappeared and later reappeared. **(D)** Mean lifetime (± S.E.M.) of DHFR^Cy5-TMP^ spots in the control experiments was substantially larger than mean lifetime of Spt5^DHFR-Cy5^ in the transcription experiments. This suggests that Spt5^DHFR-Cy5^ spot disappearance predominantly reflects dissociation of Spt4/5 from the surface-tethered transcription complex rather than photobleaching, blinking, or dissociation of Cy5-TMP from the DHFR tag. **(E)** Disappearance rate (± S.E.) of fluorescence spots (i.e., reciprocal of mean spot lifetime) in Spt5^DHFR-Cy5^ extract transcription experiments conducted at different time-fraction exposures to the 633 nm laser. No dependence of disappearance rate on exposure is observed, confirming that dye photobleaching does not significantly contribute to Spt5^DHFR-Cy5^ spot disappearance. The same laser power, with fractional exposure ≤0.9, was used in all of the transcription experiments in which DHFR-tagged proteins were observed.

**Figure S7:**
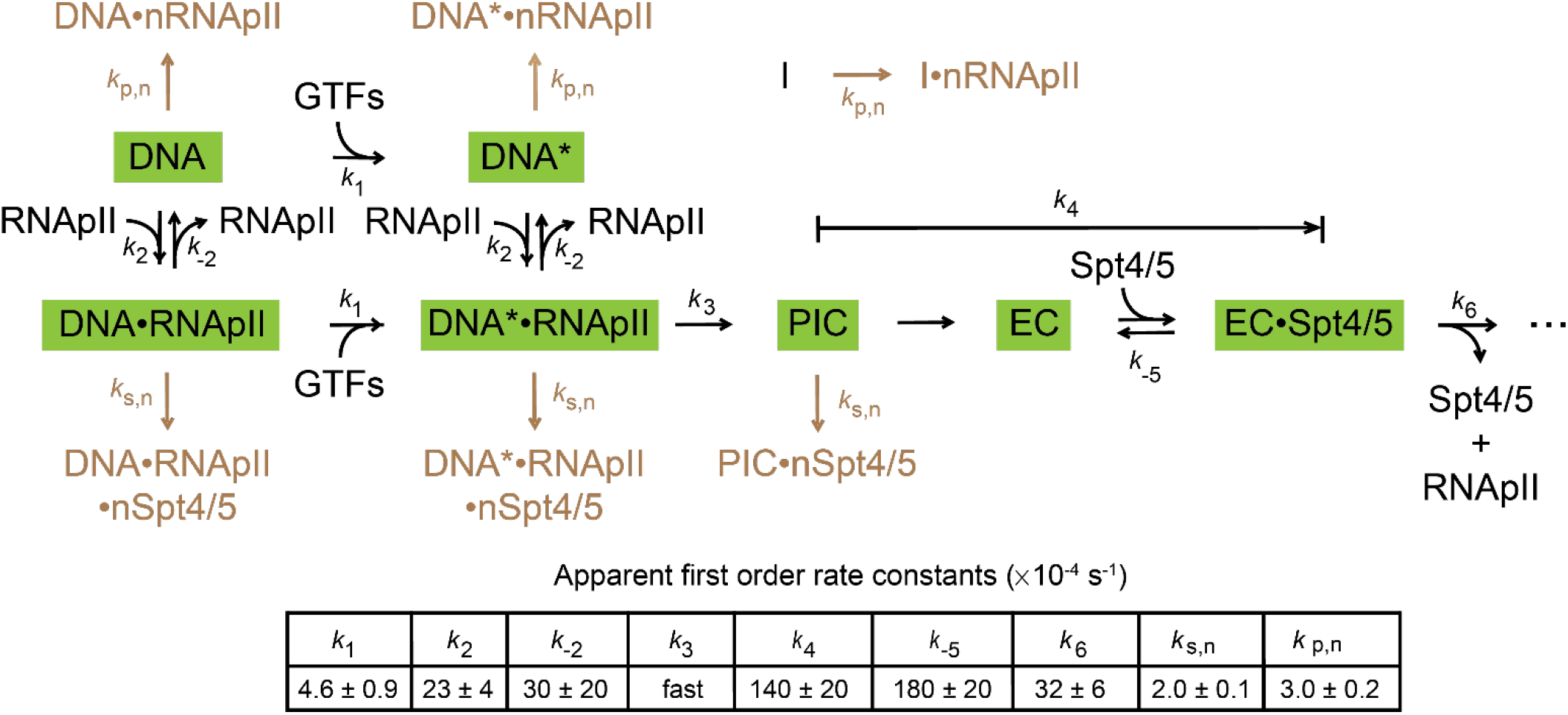
Complete scheme used in the Fig. 4 kinetic analysis. The scheme includes all species (green rectangles) and rate constants shown in **Fig. 4E** but additionally shows kinetic processes (brown) resulting in non-specifically bound (denoted with an “n” prefix) RNApII and Spt4/5. The rates of these processes (*k*_p,n_ and *k*_s,n_, respectively) were independently measured in the same experiments by analysis of randomly selected surface locations with no DNA^488^. As in previous studies (59), we deduced that some DNA^488^ spots contained molecules (“I”) incapable of protein binding; the fraction of binding-capable spots was an additional parameter determined by the global fit to be *A*_f_ = 0.62 ± 0.05.

## Supplementary Tables

**Table S1:** Kinetics for labeled Rpb1 association with DNA*.

| Extract | Condition | $A_f$ | $k_{on} (\times 10^{-3} s^{-1})$ | $k_{p,n} (\times 10^{-3} s^{-1})$ |
| --- | --- | --- | --- | --- |
| Rpb1<sup>DHFR-Cy5</sup> | +NTPs<br>+Gal4-VP16 | $0.41 \pm 0.03$ | $17 \pm 0.3$<br>($N = 212$) | $0.072 \pm 0.007$<br>($N = 117$) |
| Rpb1<sup>DHFR-Cy5</sup> | +NTPs<br>+Gal4-VP16 | $0.11 \pm 0.08$ | $2 \pm 7$<br>($N = 220$) | $1.00 \pm 0.08$<br>($N = 229$) |
| Rpb1<sup>SNAP549</sup> | +NTPs<br>+Gal4-VP16 | $0.64 \pm 0.04$ | $1.7 \pm 0.2$<br>($N = 157$) | $0.15 \pm 0.01$<br>($N = 152$) |
| Rpb1<sup>SNAP549</sup> | +NTPs<br>+Gal4-VP16 | $0.9668 \pm 0.0001$ | $5.5000 \pm 0.0006$<br>($N = 289$) | $0.59 \pm 0.03$<br>($N = 414$) |
| Rpb1<sup>SNAP549/</sup><br>Spt4<sup>DHFR-Cy5</sup> | +NTPs<br>+Gal4-VP16 | $0.86 \pm 0.03$ | $2.3 \pm 0.2$<br>($N = 146$) | $0.10 \pm 0.01$<br>($N = 118$) |
| Rpb1<sup>SNAP549/</sup><br>Spt5<sup>DHFR-Cy5 §</sup> | +NTPs<br>+Gal4-VP16 | $0.76 \pm 0.04$ | $2.2 \pm 0.3^{\pm}$<br>($N = 282$) | $0.30 \pm 0.02$<br>($N = 234$) |
| Rpb1<sup>SNAP549/</sup><br>Spt5<sup>DHFR-Cy5 §§</sup> | –NTPs<br>+Gal4-VP16 | $0.76^{**}$ | $0.47 \pm 0.08$<br>($N = 419$) | $0.2 \pm 0.01$<br>($N = 262$) |
| Rpb1<sup>SNAP549/</sup><br>Spt5<sup>DHFR-Cy5</sup> | +NTPs<br>–Gal4-VP16 | $0.76^{**}$ | $0.03 \pm 0.02^{\pm}$<br>($N = 224$) | $0.1 \pm 0.01$<br>($N = 283$) |
| Rpb1<sup>SNAP549 §§</sup> | +NTPs<br>+Gal4-VP16<br>+α-amanitin | $0.8 \pm 0.1$ | $0.68 \pm 0.02$<br>($N = 305$) | $0.17 \pm 0.01$<br>($N = 212$) |

\* $k_{on}$ and $k_{p,n}$ are the apparent first-order rate constants for binding to DNA and for non-specific binding to the chamber surface. $A_f$ is the active fraction of DNA molecules capable of RNAPII binding. See SI Methods for description of the kinetic model. Shading marks control experiments performed under conditions that produce little or no transcription.
\*\*Active fraction for these negative controls was fixed to be that of the positive experiment.
§Fit shown in **Fig. 2C**.
§§Fits shown in **Fig. S2**.
$^{\pm}$ Value shown in **Fig. 2D**

**Table S2:**
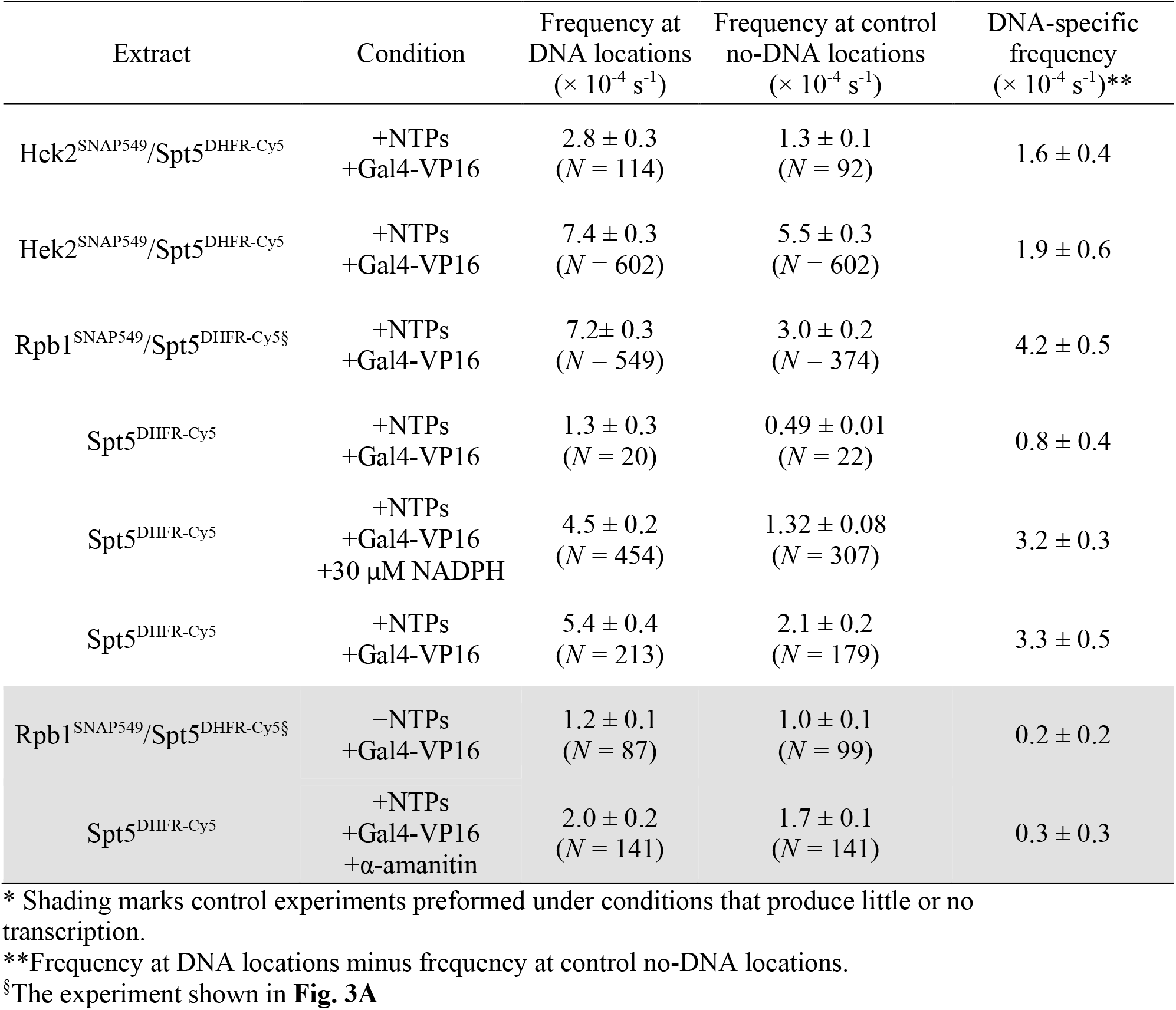
Spt5 binding frequencies*.

**Table S3:**
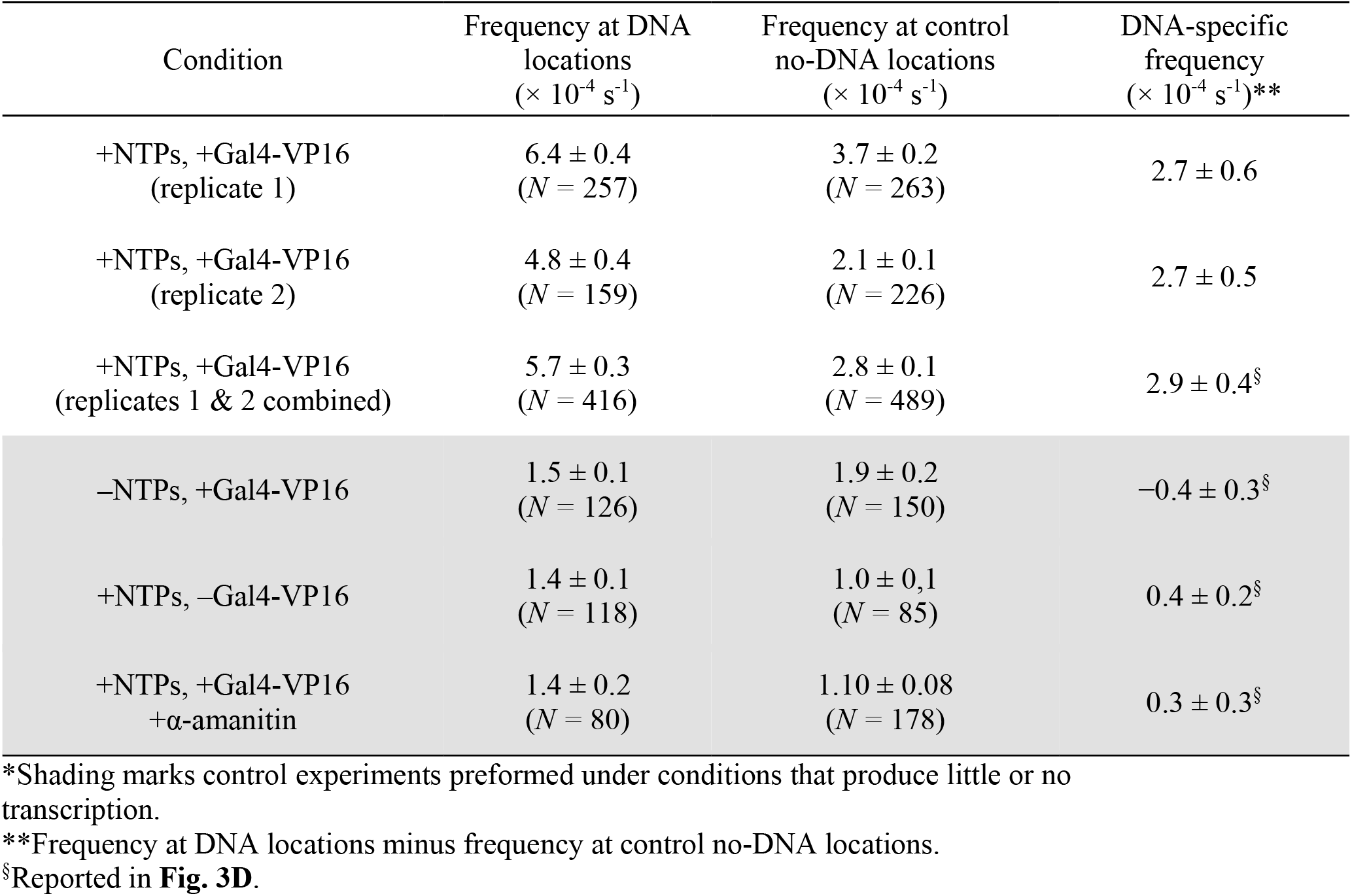
Hek2 binding frequencies in Hek2 ^SNAP549^/Spt5^DHFR-Cy5^ extract*.

**Table S4:**
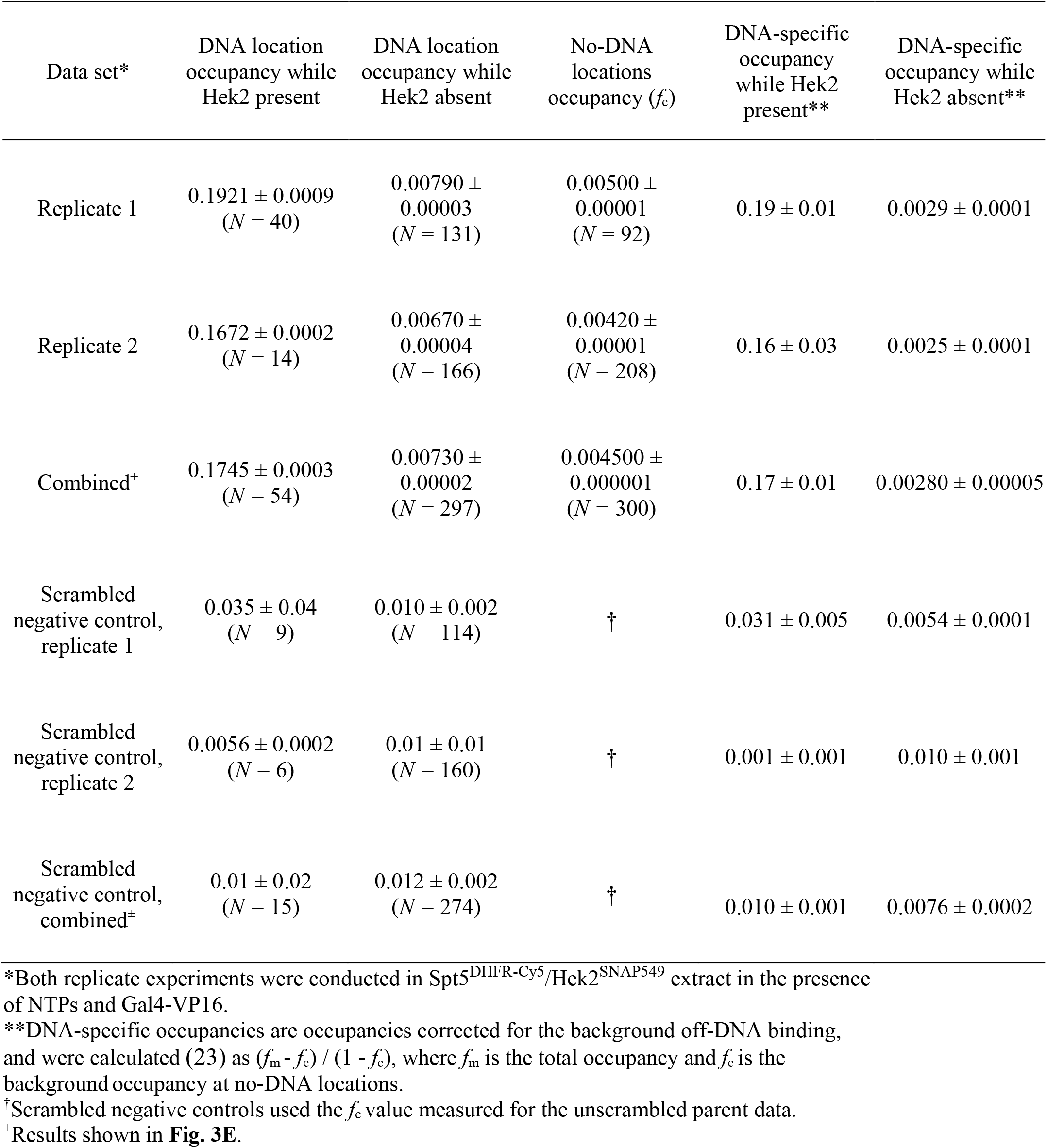
Spt5^DHFR-Cy5^ fractional occupancies on DNA molecules with/without colocalized Hek2^SNAP549^.

| Data set* | DNA location occupancy while Hek2 present | DNA location occupancy while Hek2 absent | No-DNA locations occupancy ($f_c$) | DNA-specific occupancy while Hek2 present** | DNA-specific occupancy while Hek2 absent** |
| --- | --- | --- | --- | --- | --- |
| Replicate 1 | $0.1921 \pm 0.0009$<br>($N = 40$) | $0.00790 \pm 0.00003$<br>($N = 131$) | $0.00500 \pm 0.00001$<br>($N = 92$) | $0.19 \pm 0.01$ | $0.0029 \pm 0.0001$ |
| Replicate 2 | $0.1672 \pm 0.0002$<br>($N = 14$) | $0.00670 \pm 0.00004$<br>($N = 166$) | $0.00420 \pm 0.00001$<br>($N = 208$) | $0.16 \pm 0.03$ | $0.0025 \pm 0.0001$ |
| Combined<sup>±</sup> | $0.1745 \pm 0.0003$<br>($N = 54$) | $0.00730 \pm 0.00002$<br>($N = 297$) | $0.004500 \pm 0.000001$<br>($N = 300$) | $0.17 \pm 0.01$ | $0.00280 \pm 0.00005$ |
| Scrambled negative control, replicate 1 | $0.035 \pm 0.04$<br>($N = 9$) | $0.010 \pm 0.002$<br>($N = 114$) | † | $0.031 \pm 0.005$ | $0.0054 \pm 0.0001$ |
| Scrambled negative control, replicate 2 | $0.0056 \pm 0.0002$<br>($N = 6$) | $0.01 \pm 0.01$<br>($N = 160$) | † | $0.001 \pm 0.001$ | $0.010 \pm 0.001$ |
| Scrambled negative control, combined<sup>±</sup> | $0.01 \pm 0.02$<br>($N = 15$) | $0.012 \pm 0.002$<br>($N = 274$) | † | $0.010 \pm 0.001$ | $0.0076 \pm 0.0002$ |

\*Both replicate experiments were conducted in Spt5<sup>DHFR-Cy5</sup>/Hek2<sup>SNAP549</sup> extract in the presence of NTPs and Gal4-VP16.
\*\*DNA-specific occupancies are occupancies corrected for the background off-DNA binding, and were calculated (23) as $(f_m - f_c) / (1 - f_c)$ , where $f_m$ is the total occupancy and $f_c$ is the background occupancy at no-DNA locations.
†Scrambled negative controls used the $f_c$ value measured for the unscrambled parent data.
<sup>±</sup>Results shown in **Fig. 3E**.

**Table S5:** Yeast strains and nuclear extracts.

| Strain | Genotype* | Source | Extract preparation |
| --- | --- | --- | --- |
| YF4/<br>BJ2168 | MATa, ura3-52, leu2, trp1, prb1-1122, pep4-3, prc1-407, gal2 | (69) |  |
| YF702/<br>CB012 | MATa, ura3-1, leu2-3,112, trp1-1, his3-11,15, ade2-1, pep4Δ::HIS3, prbΔ::his3, prc1Δ::hisG | (70) | Wild-type |
| YSB3161 | MATa, ura3-52, leu2, trp1, prb1-1122, pep4-3, prc1-407, gal2, SPT5-DHFR(L28C)::Hygromycin<sup>R</sup> | This study | Spt5<sup>DHFR-Cy5</sup> |
| YSB3163 | MATa, ura3-1, leu2-3,112, trp1-1, his3-11,15, ade2-1, pep4Δ::HIS3, prbΔ::his3, prc1Δ::hisG, SPT5-DHFR(L28C)::Hygromycin<sup>R</sup> | This study |  |
| YSB3338 | MATa, ura3-1, leu2-3,112, trp1-1, his3-11,15, ade2-1, pep4Δ::HIS3, prbΔ::his3, prc1Δ::hisG, RPB1-SNAP<sub>f</sub>::NATMX | This study | Rpb1<sup>SNAP549</sup> |
| YSB3403 | MATa, ura3-1, leu2-3,112, trp1-1, his3-11,15, ade2-1, pep4Δ::HIS3, prbΔ::his3, prc1Δ::hisG, SPT5-DHFR(L28C)::Hygromycin<sup>R</sup>, RPB1-SNAP<sub>f</sub>::NATMX | This study | Rpb1<sup>SNAP549</sup>/Spt5<sup>DHFR-Cy5</sup> |
| YSB3440 | MATa, ura3-1, leu2-3,112, trp1-1, his3-11,15, ade2-1, pep4Δ::HIS3, prbΔ::his3, prc1Δ::hisG, SPT5-DHFR(L28C)::Hygromycin<sup>R</sup>, HEK2-3XHA-SNAP<sub>f</sub>::NATMX | This study | Hek2<sup>SNAP549</sup>/Spt5<sup>DHFR-Cy5</sup> |
| YSB3337 | MATa, ura3-1, leu2-3,112, trp1-1, his3-11,15, ade2-1, pep4Δ::HIS3, prbΔ::his3, prc1Δ::hisG, RPB1-DHFR::Hygromycin<sup>R</sup> | This study | Rpb1<sup>DHFR-Cy5</sup> |

\*SNAP<sub>f</sub> is a fast-reacting variant of the SNAP tag (64).

